# Patterns of Structural Variation Define Prostate Cancer Across Disease States

**DOI:** 10.1101/2022.01.09.475586

**Authors:** Meng Zhou, Minjeong Ko, Anna C. Hoge, Kelsey Luu, Yuzhen Liu, Magdalena L. Russell, William W. Hannon, Zhenwei Zhang, Jian Carrot-Zhang, Rameen Beroukhim, Eliezer M. Van Allen, Atish D. Choudhury, Peter S. Nelson, Matthew L. Freedman, Mary-Ellen Taplin, Matthew Meyerson, Srinivas R. Viswanathan, Gavin Ha

## Abstract

The complex genomic landscape of prostate cancer evolves across disease states under therapeutic pressure directed toward inhibiting androgen receptor (*AR*) signaling. While significantly altered genes in prostate cancer have been extensively defined, there have been fewer systematic analyses of how structural variation shapes the genomic landscape of this disease across disease states. We uniformly characterized structural alterations across 278 localized and 143 metastatic prostate cancers profiled by whole genome and transcriptome sequencing. We observed distinct significantly recurrent breakpoints in localized and metastatic castration-resistant prostate cancers (mCRPC), with pervasive alterations in noncoding regions flanking the *AR, MYC*, *FOXA1*, and *LSAMP* genes enriched in mCRPC and *TMPRSS2-ERG* rearrangements enriched in localized prostate cancer. We defined nine subclasses of mCRPC based on signatures of structural variation, each associated with distinct genetic features and clinical outcomes. Our results comprehensively define patterns of structural variation in prostate cancer and identify clinically actionable subgroups based on whole genome profiling.

## INTRODUCTION

Over the past decade, genomic sequencing studies have progressively sharpened our view of the genetic landscape of prostate cancer (1). Such studies have defined key driver genes in prostate cancer and have enabled the deployment of therapeutic agents in molecularly-defined disease subsets, including potent androgen receptor (*AR*)-targeted therapies (2, 3), poly (ADP-ribose) polymerase (PARP) inhibitors in *BRCA1/2*-altered prostate cancers, and immune checkpoint inhibitors in cancers with microsatellite instability (4–7).

To date, most cancer genomic studies have employed whole exome sequencing (WES) and have thus been focused on mutations or copy number alterations that occur within the protein-coding regions of genes, which represent only 1-2% of the genome. More recent studies applying whole genome sequencing (WGS) to prostate and other cancers have identified previously underappreciated recurrent alterations in regulatory (non-coding) regions of the genome and have illuminated complex mechanisms of genomic alterations – driven by structural variants (SVs) – that are difficult to discern by WES; in the case of prostate cancer, most of these studies have focused on localized disease, the disease state in which tissue is most readily accessible for profiling (8–22). There remains a need for continued high-resolution genomic discovery efforts in prostate cancer.

In addition to efforts characterizing entire cancer genomes, recent studies have illustrated the importance of molecularly profiling prostate cancer across disease states. While many localized prostate cancers can be cured with surgery or radiotherapy, a substantial portion of higher-risk cancers recur and progress to metastatic disease, which is incurable. Recurrent prostate cancer may have a long natural history, during which time a patient may receive several lines of therapy – with androgen deprivation therapy (ADT) as a backbone – that may shape the cancer’s genomic landscape (23).

Indeed, while hormone-refractory castration-resistant prostate cancer (CRPC) has been less extensively profiled than localized prostate cancer, several studies have indicated that CRPCs display genomic landscapes distinct from treatment-naïve disease (24, 25). A cardinal hallmark of CRPC is the reactivation of *AR* signaling in the face of maximal ADT (22, 26–28). This may occur via diverse mechanisms, including the production of constitutively active *AR* splice variants (*AR-V*s) and activating mutations or copy number amplifications of the *AR* gene (29–31) or of regulatory elements distal to the gene body (13, 15, 32). To date, the relative contribution of each of these mechanisms in driving *AR* reactivation in CRPC has not been systematically explored. Also needed is a more global map of significant hotspots of structural variation in prostate cancer genomes, drawn within a rigorous statistical framework.

In this study, we performed linked-read WGS on 36 mCRPC tumor-normal pairs. We combined these data with WGS and whole transcriptome sequencing (RNA-Seq) data from previously described localized and metastatic CRPC cohorts (9, 13, 15, 33). We then established a harmonized workflow for the integrative genomic analysis of 278 localized and 143 metastatic CRPC samples, interrogated both hotspots and genome-wide patterns of structural variation, and evaluated their consequences.

## RESULTS

### WGS analysis of localized and metastatic prostate cancer cohorts

We performed linked-read whole genome sequencing on 36 biopsy specimens from 33 mCRPC patients and matched blood normal controls. After quality control, 17 tumor samples were excluded based on insufficient tumor purity and/or contamination, reflecting the challenge of obtaining high-purity metastatic biopsies, particularly from bone lesions (34). We included only samples with tumor purity > 15% in downstream analyses so as to increase confidence in structural variant calls (**Methods,** **Figure 1A, Table S1**). We re-analyzed a linked-read WGS dataset of 23 samples published previously (15), resulting in a total of 42 linked-read WGS samples from 38 patients with mean coverage of 34X (range 21X - 54X) and 33X (range 25X - 45X) for tumor and normal samples, respectively (**Table S1A**). The mean molecule length was 29 kB and 34 kB in tumor and normal samples, respectively (**Table S1A**).

**Figure 1.**
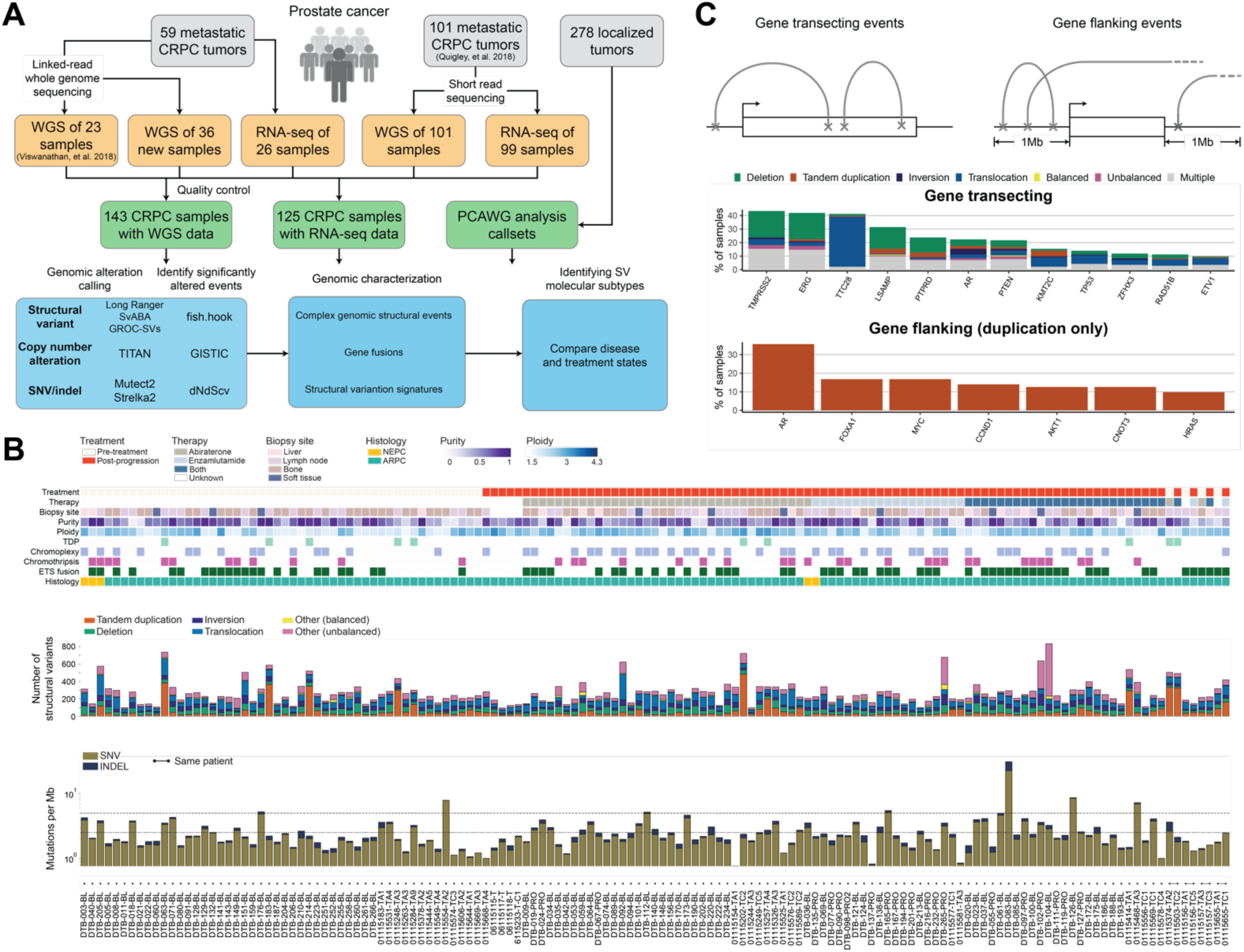
Study design and genomic landscape of mCRPC. **(A)** Workflow of study and data analysis. Tumor specimens (grey) from both primary prostate cancer and mCRPC were included in this study. Linked-read and short-read whole-genome sequencing (WGS) and RNA-sequencing datasets were either generated for this study or reanalyzed from prior studies (13, 15). A pooled dataset of 143 mCRPC samples with WGS data was used in this study after curation (**Methods**). Genomic alteration call-sets for 278 primary localized prostate cancer samples were obtained from ICGC/TCGA Pan-Cancer Analysis of Whole Genomes (PCAWG) (9, 33). For 125 mCRPC samples, RNA-seq was used. The overview of the genomic alteration and characterization analysis is shown. **(B)** Clinical annotations and somatic alterations for 143 patient samples in the pooled mCRPC cohort. Samples are ordered by treatment type; the four patients with pre-treatment and post-progression pairs are placed at the right. (Top) Clinical and sample information and genomic pattern classifications. (Middle) Distribution of genomic rearrangement types in individual samples. (Bottom) Mutational burden for SNVs and indels computed as number of mutations per mega-base pair (Mb). Y-axis shown in logarithmic scale. Threshold lines indicates mutational burden at 2.5 and 5 mutations per Mb. **(C)** Genomic rearrangement alteration profiles of key mCRPC genes. (Top) Events were categorized into gene transecting and gene flanking events (**Methods**). Gene transecting: if any of its breakpoints was located within the gene body region. Gene flanking: rearrangements which were not gene transecting and had breakpoints located within 1 Mb of either transcription start site or termination site of the gene. Only 159 genes reported and known to be involved in prostate cancer were considered in this analysis (**Table S1G and S1H**). (Middle) Frequency and distribution of rearrangement types for gene transecting events; genes with ≥ 10% frequency are shown. Gene transecting events were prioritized over flanking events during annotation. The category “Multiple” represents gene-sample pairs carrying more than one type of rearrangement event. (Bottom) Frequency of gene flanking events by tandem duplication; genes with ≥ 10% are shown.

We further combined these data with 101 mCRPC samples sequenced with standard short-read sequencing, published previously (13). This resulted in the generation of a final combined cohort of 143 tumor-normal pairs, which were uniformly analyzed for copy number and structural alterations via a harmonized pipeline (**Figure 1A**). Fifty-four samples (37.8% of 143 samples) were collected at castration resistance, prior to receiving treatment of second-generation androgen receptor signaling inhibitor (ARSi) such as abiraterone and/or enzalutamide (“pre-treatment”), while the remaining 89 samples (62.2% of 143 samples) were collected at progression (“post-treatment”, **Figure 1B**, **Table S1B**). We analyzed the somatic single nucleotide variant (SNVs), insertion-deletions (indels), copy number alterations (CNAs), and SVs in the combined cohort and identified recurrent somatic alterations in each of these classes (**Figure 1A, Methods**).

A total of 2,315,452 SNVs and indels were called, with a mean tumor mutation burden (TMB) of 2.82 mutations per million bases (Mb). We confirmed that known driver genes of prostate cancer were enriched for non-synonymous mutations, including *TP53*, *RB1*, *PTEN*, *FOXA1*, *CDK12*, *AR* and *SPOP* among known COSMIC Cancer Gene Census genes (dndscv, q ≤ 0.1, **Table S1C and S1D, Methods**). We detected an average of 272 (range 96-833) SV events per sample. Based on breakpoint orientations, SV events were classified into deletions, inversions, tandem duplications, inter-chromosomal translocations, and intra-chromosomal translocations, while intra-chromosomal translocations were further divided into balanced and unbalanced events based on copy number information (**Methods**). Chromoplexy was detected in 53 samples (37.1% of 143 samples) while chromothripsis was detected in 37 samples (25.9%); these events were not mutually exclusive (Fisher’s exact test, log-odds=1.417, p-value=0.612). Ten cases (7.0%) harbored a genome-wide tandem duplicator phenotype (TDP), all of which had *CDK12* inactivating alterations, as recently reported (15, 35). We found that TDP was mutually exclusive with ETS rearrangements (Fisher’s exact test, log-odds ratio=0.133, p=0.043) and chromothripsis (log-odds ratio=0.301, p-value=0.007), as previously reported (10, 13, 15, 35).

Analysis of CNA events across the genome revealed amplification and deletion peaks in the regions of known prostate cancer genes (10, 13, 15, 24). Many oncogenic drivers of mCRPC, such as *AR* and *MYC*, are within peaks of amplification across the cohort, while tumor suppressors such as *PTEN, TP53,* and *KMT2C* were found within deletion peaks (**Figure S1C**, **Table S1E and S1F**), consistent with prior reports (10, 13, 15, 36).

### Recurrent somatic structural variants in prostate cancer-associated genes

Structural variants may either activate or inactivate gene function, depending on the location of the breakpoints and the specific class of SV. We analyzed the potential impact of SVs called across our combined cohort, distinguishing between those with predicted inactivating (“gene transecting events”) and activating (“gene flanking events”) effects (**Figure 1C, Figure S1C, Table S1G and S1H**). Frequent gene transecting alterations were observed at the *TTC28* (37.1% of 143 samples), *LSAMP* (31.5%), and *PTPRD* (23.8%) loci, which have not been extensively studied in prostate cancer, though they have been reported in callsets for certain cohorts (28). Rearrangements involving *TTC28* were predominantly inter-chromosomal translocations between the gene body and various non-recurrent partner loci (**Figure S2E**). This likely represents retrotransposon activity, given that the *TTC28* locus harbors an active L1 retrotransposon element (37–39). Transecting SVs within the *LSAMP* and *PTPRD* genes were predominantly deletions. Both of these genes are sites of deletion/rearrangement in cancer and have been reported to function as tumor suppressors, though they have not been extensively studied within the context of prostate cancer (40–43) (**Figure 1C**). Of note, although gene transecting events would be predicted to disrupt individual genes, the most frequent transecting events identified via this analysis were deletion events that span the adjacent *TMPRSS2* and *ERG* genes (observed in 37.8%), which actually produces an activating*TMPRSS2-ERG* fusion.

Duplication events that flank an intact gene could activate oncogenes, either by resulting in copy number gain of the gene or by duplicating non-coding regulatory regions (13, 15). In our combined cohort, we observed recurrent tandem duplication events with breakpoints located in the flanking gene regions of several known prostate cancer oncogenes, including *AR* (35.7%), *FOXA1* (16.8%), *MYC* (16.8%), and *CCND1* (14.0%), consistent with frequencies that have been previously reported by us and others (10, 13, 15) (**Figure 1C**).

Certain prostate cancer driver genes were altered by multiple classes of structural alterations in both the gene body and flanking regions (e.g., *AR*, *PTEN*), while others were predominantly altered by a single alteration class (e.g., SNVs for *TP53*, intragenic translocations for *TTC28*, or flanking tandem duplications for *MYC*) (**Figure 1C, Figure S1C**). Collectively, these results catalog how diverse classes of rearrangements, both within genes and in intergenic regions, alter prostate cancer genes across disease states.

### Significantly recurrent breakpoint regions in the mCRPC genome are enriched within enhancer regions and *AR* binding sites

Next, we sought to identify significantly recurrent breakpoint (SRB) regions across our combined mCRPC cohort of 143 cases in a genome-wide, unbiased manner. We applied a Gamma-Poisson regression approach to model the occurrences of SV breakpoints within 100 kB windows across the cohort as previously described (44). Importantly, this model nominates significantly recurrent breakpoint regions likely to function as cancer drivers by accounting for six different covariates, including sequence features (e.g., GC-content and transposable elements), fragile sites, heterochromatin regions, DNase I hypersensitivity sites (DHS), and replication timing (**Methods**), which may increase specificity over prior studies that have accounted only for SV frequency or for breakpoint density within a genomic window (10, 13, 15, 45).

We identified a total of 55 significantly recurrent breakpoint regions genome-wide across our combined mCRPC cohort (Benjamini-Hochberg corrected, q-value ≤ 0.1, **Figure 2A**, **Table S2A**). Thirty-six (65.5%) SRB regions were located within 1 Mb of 14 known prostate cancer driver genes, including *AR* and its enhancer, *TMPRSS2/ERG*, *TP53*, *PTEN*, *FOXA1*, and *MYC*. For these 14 driver genes, we did not observe significant differences in SV alteration frequencies when comparing between pre-treatment (N=54) and post-progression (N=89) samples, except in the case of *ERG, for* which the SV frequency was enriched in pre-treatment samples (Fisher’s exact test, p = 0.0395; all other genes had p > 0.05, **Figure S3B**). We also did not identify any major differences in the alteration frequencies of prostate cancer genes in four patients who had paired samples collected both before treatment with and after progression on an ARSi. (**Figure S3A**).

**Figure 2.**
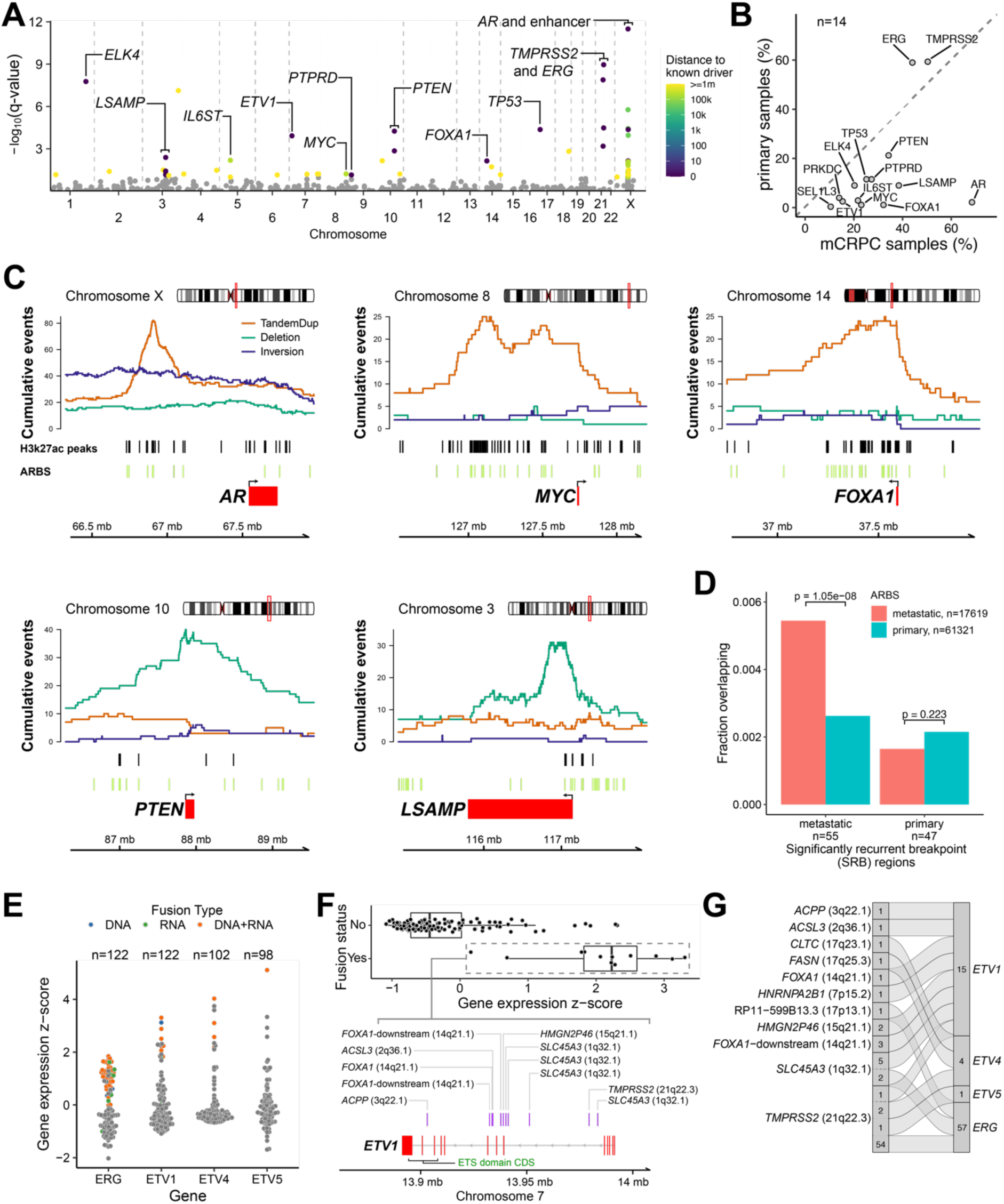
Genome-wide analysis of genomic rearrangements in mCRPC. **(A)** Analysis of significantly recurrent breakpoint (SRB) identified regions of rearrangement hotspots, genome-wide, using a Gamma-Poisson regression model. Each dot corresponds to a 100 kB bin (n=26,663 total bins). Statistically significant SRB bins with FDR (Benjamini-Hochberg) q-value ≤ 0.1 (n=55) are colored based on the distance to the nearest known prostate cancer driver gene, within 1 Mb. The driver genes within 1 Mb of the SRB bins are labeled. A square bracket is used for genes spanning multiple bins. Bins with q-value > 0.1 were not significant (grey). **(B)** Comparison of SV alteration frequency in mCRPC versus primary localized prostate cancer. The union set of genes (n=14) within 1 Mb of SRB hotspot regions in mCRPC and localized prostate cancer cohorts was included in the comparison. The frequencies represent total gene transecting and flanking SV events. All labeled genes were significantly enriched in either mCRPC or primary localized tumors (Fisher’s test, p-value < 0.05). **(C)** Patterns of rearrangements at the loci of driver genes identified at SRB regions in mCRPC cohort of 143 tumors. Cumulative counts of intra-chromosomal SV events (tandem duplications “TandemDup”, deletions, and inversions) were computed based on the breakpoints and span of the events. Histone H3 lysine 27 acetylation (H3K27ac) and *AR* binding sites (ARBS) specific to mCRPC were obtained from a previous study (46). Inter-chromosomal translocations are not shown. Genome coordinates based on hg38 build. **(D)** Overlap of *AR* binding sites (ARBS) within SRB hotspots of mCRPC (55 regions) and primary localized prostate (47 regions) cohorts. Metastatic-specific and primary localized-specific ARBS were obtained from previous studies (46, 85). χ^2^ test of independence p-values are shown. **(E)** Fusion status and expression of selected genes in ETS transcription factor gene family in the mCRPC cohort with WGS and RNA-seq data. Fusion type was defined as the data evidence that supported the event: DNA-only, corresponds to WGS; RNA-only, corresponds to RNA-seq; DNA+RNA, corresponds to support from both WGS and RNA-seq. Each dot represents a tumor sample and is colored based on fusion type of each sample; grey indicates no evidence of fusion event. Data shown for samples with available expression data for the specific ETS gene. Gene expression values of full-length transcripts are z-score normalized. **(F)** Fusion profile of *ETV1*. DNA rearrangement breakpoints supporting the fusion (purple bars) are indicated with the corresponding fusion partners. Exons of the ETS domain (red) are indicated. Genome coordinates based on hg38 build. **(G)** Summary of fusion partners for selected genes in ETS transcription factor gene family in mCRPC cohort. Fusion events and partners are indicated by flow connections. Total counts of individual fusion events and partners across the cohort are shown.

We then sought to compare how SVs drive prostate cancer across disease states. For the localized disease state, we utilized genome alteration calls from 278 primary localized prostate cancer tumors from the PCAWG study (9, 33). Using Gamma-Poisson regression, we first identified 47 SRB regions in localized prostate cancer tumors (**Figure S2B, Table S2B**). Six prostate cancer genes (*TMPRSS2*, *ERG*, *TP53*, *PTEN*, *IL6ST*, *ELK4*) within mCRPC SRB regions were also found within or in proximity (less than 1 Mb) to an SRB region in localized disease. By contrast, four SRBs (three near *SEL1L3* and one near *PRKDC*) were unique to localized disease, while 27 SRBs were unique to mCRPC with six genes nearby (*LSAMP*, *ETV1*, *MYC*, *PTPRD*, *FOXA1*, *AR*). When comparing SV alteration frequencies for the 14 genes located within SRB regions in either mCRPC or localized tumors, 12 genes were significantly more altered in mCRPC samples, while *TMPRSS2* and *ERG* were significantly more altered in localized disease (Fisher’s exact test, p < 0.05 for all genes, **Figure 2B**). We repeated this comparison using a non-overlapping cohort of localized prostate cancers profiled by WGS and found similar genes enriched for SVs in either the localized or metastatic disease states (**Figure S2C-E**) (18). Thus, localized prostate cancer and mCRPC have significantly different landscapes of recurrent SVs.

To explore the potential functional consequences of SVs in intergenic SRB regions, we overlapped SV breakpoints with locations of H3K27ac marks specific to mCRPC (46). We observed that intergenic SVs within SRB regions in the mCRPC cohort included gene flanking events that were enriched at putative enhancer regions for *AR*, *MYC*, and *FOXA1*, which all had frequent focal duplication events at sites marked by mCRPC-specific H3K27ac deposition (**Figure 2C, S2F**). Interestingly, an intragenic deletion SRB region was observed near the transcription start site of *LSAMP*, also overlapping H3K27ac marks. *PTEN* had a high level of both gene transecting and flanking deletions, leading to SV breakpoints that were spread more broadly around the gene.

We also observed an enrichment of metastatic-specific *AR* binding sites (ARBS) compared to localized primary ARBS within the 55 mCRPC SRB regions (**Figure 2D**, one-sided proportion test, p = 1.05 x 10^-8^). This enrichment was not observed for localized primary SRB regions (p = 0.22). These results highlight that SVs within mCRPC SRB regions may be capturing the genome-wide footprint of activated *AR* signaling that occurs with castration resistance.

### Refined landscape of ETS gene fusions from integrated analysis of the genome and transcriptome

We applied gene fusion analysis by integrating both genome rearrangements and fusion RNA transcript information from 127 samples with RNA-seq data (**Figure 1A**, **Table S2C**, **Methods**). For gene fusions involving E26 transformation-specific (ETS) transcription factor gene family members (*ERG*, *ETV1*, *ETV4* and *ETV5*), we detected 50 events supported by both DNA and RNA evidence, 15 supported by only DNA evidence, and 10 supported by only RNA evidence (**Figure 2E**, **Figure S2G**). Overall, 74 samples (51.7% of 143 samples) harbored a fusion event of the *ETS* gene family, consistent with previous reports (47, 48) (**Figure 1B, Table S2C**).

Among the ETS fusions, *ERG* was most commonly involved with *TMPRSS2* as the fusion partner (54 out of 57 cases, **Figure 2G**). Other common ETS fusion partners were *SLC45A3* (7 cases) and lncRNA RP11-356O9.1 downstream of *FOXA1* (3 cases). *ETV1* had eight distinct fusion partners, which is consistent with previous reports that *ETV1* is a promiscuous ETS fusion member (49) (**Figure 2F**).

We observed that fusions of the ETS family members *ERG*, *ETV1*, *ETV4* and *ETV5* were mutually exclusive, except for one sample which harbored fusions of both *ERG* and *ETV1* (**Figure S2G**). In addition, gene fusion events were correlated with higher expression of the corresponding ETS genes they involved (Wilcoxon rank-sum tests, p < 0.05 for all genes, **Figure 2E**). In the 38 cases which did not show any evidence for an ETS fusion, we noted that presence of high-level expression (z-score > 1) of ETS genes *ERG*, *ETV1*, *ETV4*, and *ETV5* were also mutually exclusive (Fisher’s exact test, p = 0.480 for *ETV4*, p = 0.363 for *ETV5*, **Figure S2G**). These may represent cases of missed fusion calls, or cases in which ETS family members are transcriptionally activated through non-genetic mechanisms.

Interestingly, we also observed 20 cases (14.0% of 143 cases) involving fusions between the ETS family member *ELK4* and its upstream gene *SLC45A3*. While the *ELK4* locus was an SRB in our analysis (**Figure 2A** and **Figure S2G**), manual inspection of individual samples revealed evidence for a genomic event capable of producing an *ELK4* fusion in only 1 out of 20 cases (**Figure S2G and data not shown).** In contrast, 19 other cases showed *ELK4* fusions on RNA-sequencing alone, consistent with a mechanism of cis-splicing or transcriptional read-through events that may perhaps be induced by local genomic alterations (50–52) (**Table S2C**). Importantly, although *ELK4* fusions were significantly correlated with higher expression of *ELK4* (Wilcoxon rank-sum test, p = 7.91x10^-5^, **Figure S2G**), these events were not mutually exclusive with fusions of other ETS family members (Fisher’s exact test, p = 0.472). Thus, the functional consequences of these *ELK4* fusions and whether they contribute to prostate cancer pathogenesis in a manner similar to other ETS fusions remains to be determined.

### Classes of rearrangements driving *AR* signaling in mCRPC

Genomic alterations involving the *AR* locus play an important role in sustaining *AR* signaling in mCPRC (13, 15, 26, 53). We sought to catalog the spectrum of diverse structural mechanisms that underlie *AR* activation in mCRPC, and the relationship between them, in our combined mCRPC cohort. To understand the relationship between different modes of somatic *AR* activation, we first determined copy number at the *AR* gene body and its upstream enhancer and categorized samples into distinct groups of: (**1**) co-amplification (N = 99, 69.2% of 143 cases); (**2**) selective *AR* gene body amplification (N = 4, 2.8% of 143 cases); (**3**) selective *AR* enhancer gains (N = 17, 11.9% of 143 cases), and (**4**) lack of amplification for both (N = 23, 16.0% of 143 cases) (**Figure 3A-C, Table S3**). For the 122 samples with expression data available, we observed that *AR* gene expression was higher in the co-amplification and selective enhancer categories compared to samples with no amplification, after accounting for tumor purity and ploidy (ANCOVA/TukeyHSD p-values 5.6x10^-11^ and 4.5x10^-4^, respectively), but not for selective *AR* status (ANCOVA p = 0.098) (**Figure 3B, Methods**). Interestingly, we observed that samples with selective enhancer duplication exhibited similar *AR* expression levels to samples with co-amplification (ANCOVA, p = 0.31), even though enhancer duplications involved lower-copy gains (mean 2.73, range 1.97 - 5.02) compared to co-amplified samples (mean 12.87, range 1.55 - 150.57) (**Figure 3A**). This is consistent with previous results (15, 28) and suggests a mechanism whereby *AR* expression levels are increased through even modest genomic expansion of enhancer elements.

**Figure 3.**
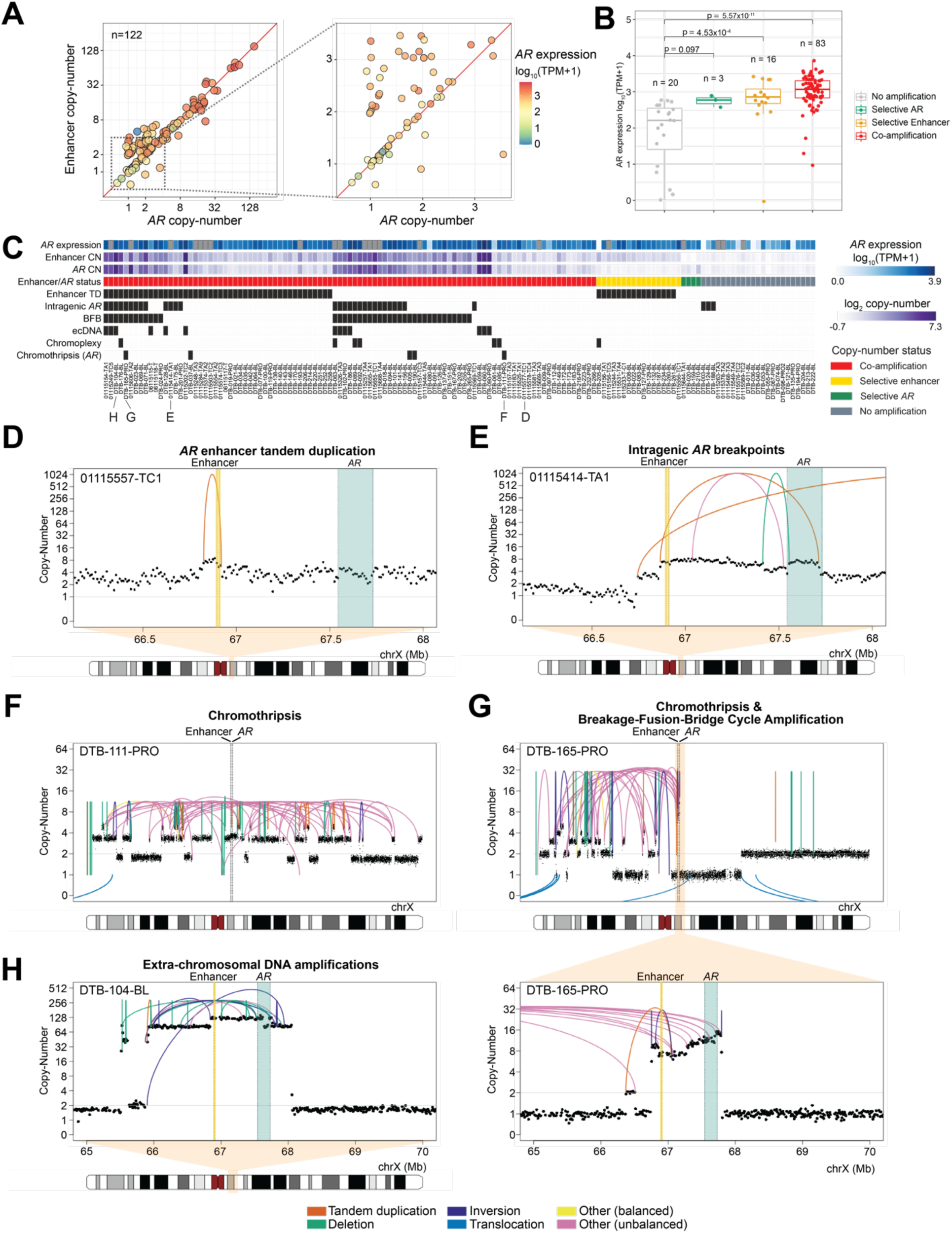
Modes of *AR* activation in mCRPC. **(A)** Copy number of *AR* gene and its enhancer (∼624 kB upstream) for mCRPC cohort samples after adjustment by tumor purity and sample ploidy normalization. Data shown for samples with available *AR* gene expression data. (Left) Copy number of *AR* and its enhancer are shown in log_2_ scale, colored based on *AR* gene expression level (transcripts per million, TPM). (Right) Excerpt of figure highlighting *AR* expression for samples with lower copy number values. **(B)** *AR* expression for *AR* locus copy number status for 122 samples with available *AR* gene expression data. ANCOVA test was performed to account for tumor purity and ploidy as covariates. TukeyHSD p-values for pair-wise comparisons between groups with *AR* locus amplification status and groups with no amplification. **(C)** Patterns of rearrangements involving the *AR* locus in 143 mCRPC samples. Presence of specific alteration events and complex rearrangements (black) were predicted automatically and manually curated. *AR* gene expression shown (blue shades) for same samples in (B); samples with no available expression data are indicated in grey. Representative examples of each category are presented in (D) to (H). **(D-H)** Complex and simple rearrangement patterns involving the *AR* locus, including focal duplication events on *AR* enhancer **(D)**, intragenic amplification event leading to a breakpoint within *AR* between exons 4 and 5 **(E)**, chromosomal level chromothripsis events involving *AR* and enhancer **(F)**, arm-level chromothripsis coinciding with *AR* amplification by break-fusion-break cycle **(G)**, extra-chromosomal DNA amplicon including *AR* and enhancer **(H)**. *AR* gene boundary (green) and its enhancer (yellow) are shown; concave arcs, intra-chromosomal SV events; convex arcs, inter-chromosomal SV events. Copy number values represent 10 kB bins and have been tumor purity corrected.

We then systematically and manually curated the diverse mechanisms of rearrangements activating *AR* signaling by analyzing patterns of SVs at the *AR* locus (**Figure 3C, Table S3, Methods**). We observed a total of 62 samples (43.4% of 143 samples) with tandem duplication SV events that spanned the enhancer with breakpoints located within 1 Mb, including 16 cases (11.2% of 143 samples) with selective enhancer copy number amplification status (**Figure 3D**). Thirty-two samples (22.4% of 143 samples) harbored intragenic rearrangements within *AR*, which may have implications for the production of truncated, constitutively-active *AR* splice variants (31). For example, in case 01115414-TA1, we observed a tandem duplication breakpoint selectively amplifying exons 1-4 of AR, but not exons 5-8 of *AR*, which includes the ligand binding domain; such an event could promote selective expression of a constitutively active truncated *AR* variant, although RNA-Seq data was not available on this sample. (**Figure 3E**). In another case, DTB-124-BL, our reanalysis confirmed that a focal deletion involving exons 1-4 resulted in the expression of truncated *AR* variants as previously described (28) (**Figure S3C**). Interestingly, in the 21 samples with selective *AR* enhancer or selective *AR* gene body copy number gain, none harbored intragenic SV events in *AR*.

We also examined the landscape of complex rearrangement mechanisms involving *AR*; these mechanisms involve multiple SV events and copy number patterns, including chromothripsis, extrachromosomal DNA (ecDNA), chromoplexy, and breakage-fusion-bridge cycle (BFB) (**Methods**). Chromothripsis of a region or the entire X chromosome involving the *AR* locus was detected in 5 samples, all of which had co-amplification of *AR* and enhancer, suggesting that following repair after catastrophic DNA shattering the *AR* locus was retained or further amplified (**Figure 3F**, **Figure 3G**). Thirteen samples (9.1% of 143 samples) showed very high levels of *AR* and enhancer copy number, suggesting the possibility of their presence on extrachromosomal elements (ecDNA, **Figure 3H**). In 40 samples (28.0% of 143 samples), the most frequent complex rearrangement mechanism, BFB, led to *AR* locus amplification, including instances following chromothripsis (14, 54) (**Figure 3G**). Overall, we noted that complex rearrangement events, which frequently co-occurred, were significantly enriched in samples with co-amplification of *AR* and enhancer compared to those with selective enhancer copy number gain status (Fisher’s exact test, p = 1.52x10^-4^).

### Distinct signatures of structural rearrangement patterns in mCRPC

To systematically characterize genome-wide structural rearrangement patterns in mCRPC, we performed rearrangement signature analysis using SV breakpoint features, non-negative matrix factorization, and known reference signatures (12, 55) (**Methods**). First, we derived signatures *de novo*, which identified eight signatures: six that matched reference signatures (RefSigs) also observed in localized prostate cancer (> 0.91 cosine similarity), one that matched an ovarian cancer RefSig.R14 associated with large segment (100 kB-10 Mb) TDP (0.96 cosine similarity), and one that was likely an artifact specific to linked-read sequencing (**Figure S4A-C, Table S4A and 4B**). Therefore, we excluded the linked-read data and focused on standard WGS data from 101 mCRPC cases for further SV signature analysis. We fit these samples to the nine known RefSigs from localized prostate cancer (R1-4, R6a-b, R8-9, R15) and the one (R14) from ovarian cancer (**Figure S4A**, **Table S4C**). Overall, eight of the RefSigs were detected across our cohort (R1-2, R4, R6a-b, R9, R14-15). Notably absent in mCRPC were RefSig.R8 (short, 1-10 kB inversions) and RefSig.R3, which is associated with germline *BRCA1* mutations and short (1-100 kB) tandem duplications (11, 12, 55, 56) (**Figure S4D**). By contrast, we observed increased prevalence of some signatures in mCRPC compared to localized disease, including RefSig.R2 (large SV classes, abundant translocations; 97% vs. 60%), RefSig.R4 (clustered translocation events; 37% vs. 27%), and RefSig.R15 (large deletions and inversions, 48% vs. 37%) (**Figure S4D**).

To investigate whether molecular subtypes in mCRPC can be grouped based on SV patterns, we applied hierarchical clustering on the exposure of the eight fitted signatures and identified nine distinct SV clusters (**Figure 4, Table S4C**). We observed that samples in SV Cluster 1 were composed of non-clustered translocation events and were significantly enriched for the presence of chromoplexy (χ^2^ test, FDR corrected, q = 0.12). SV Cluster 3 was characterized by many short deletions and was significantly enriched for *BRCA2* mutations (q = 5.01x10^-4^). SV Cluster 5 was significantly enriched for *SPOP* mutations (q = 0.02), with no instances of ETS gene family fusion (q=0.06), consistent with previous reports (57). SV Cluster 6 had the highest prevalence of *TP53* mutation (q = 0.02), while SV Cluster 7 samples harbored the TDP associated with *CDK12* inactivation (q = 3.52 x 10^-11^) as well as enrichment for *CCND1* gains (q = 0.02), consistent with previous reports (35, 58). The remaining clusters did not have enrichment for any alterations in known driver genes; however, distinct SV patterns were still evident in SV Cluster 4 (non-clustered tandem duplications), 8, and 9 (increased clustered SV events of various classes).

**Figure 4.**
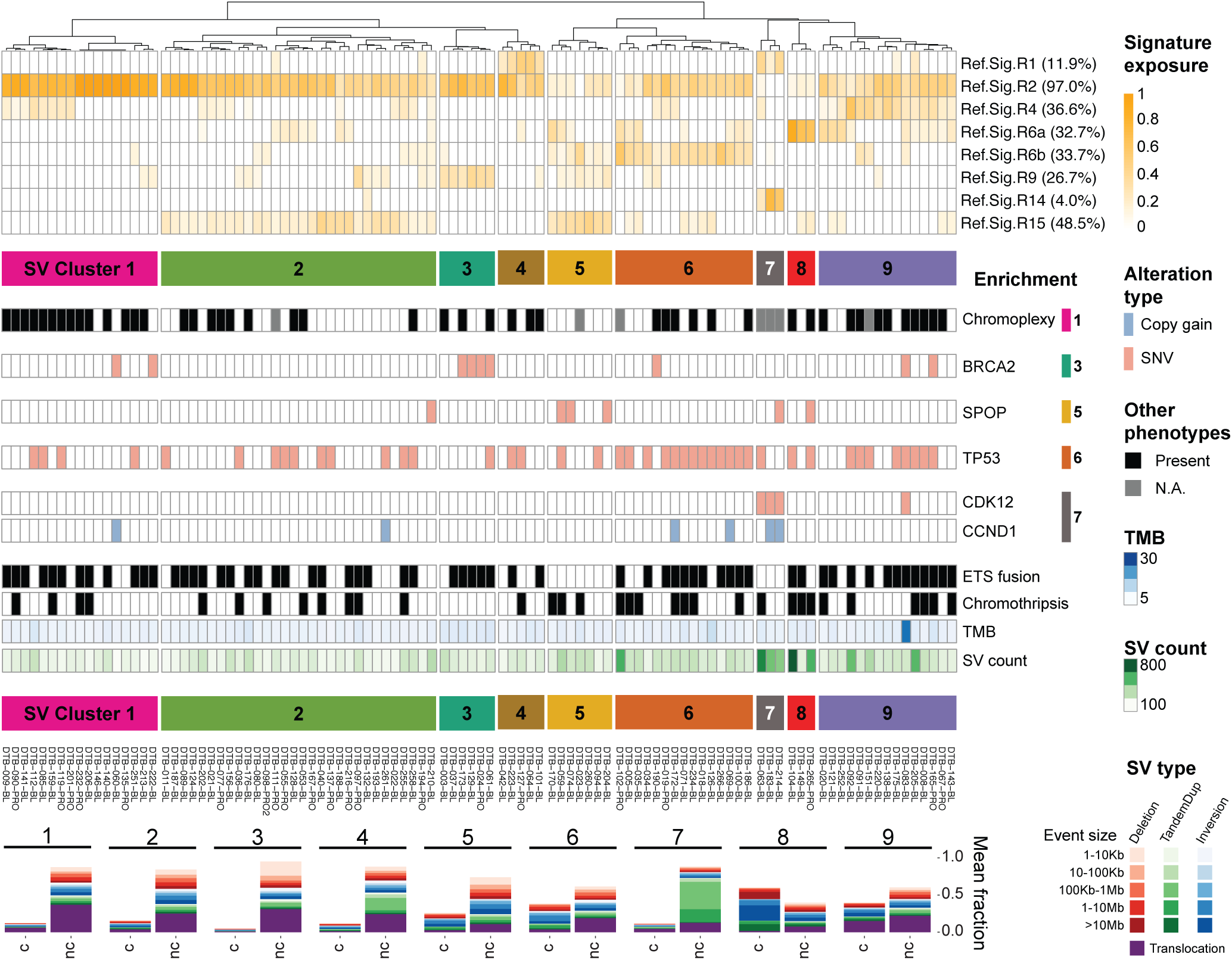
Clustering of mCRPC SV signatures. SV signature analysis and hierarchical clustering identifies nine distinct molecular groups. (Top) Dendrogram of the clustering of SV signature exposure. The prevalence of each signature was computed based on having ≥ 0.05 exposure (proportion of SVs). (Middle) Enrichment of altered prostate cancer drivers. Enriched alterations in Cluster 1, 3, 5, 6, and 7 are shown based on statistical significance by χ^2^ test. (Bottom) Composition of SV types and sizes for each SV cluster, separated by non-clustered (nc) and clustered (c) SV events.

While SV Clusters 3, 5 and 6 had significant enrichment of mutations in *BRCA2*, *SPOP,* and *TP53*, respectively, not all samples within each cluster harbored these mutations. Intriguingly, we further noted that clinical outcomes showed significantly better stratification when using SV Clusters 3, 5, and 6 for outcome stratification compared to using the associated mutation status itself (**Figure S4D-E**). Specifically, SV Cluster 5 had significantly better overall survival than SV Clusters 3 and 6 (log-rank test, p=0.01), while the sample group with *SPOP* mutations did not have significantly greater survival compared to the sample groups with *BRCA2* and *TP53* mutations (log-rank test, p=0.45) in this cohort. Together, these results indicate the analysis of genome-wide patterns of rearrangements may provide a way to further refine molecular subtypes in mCRPC.

## DISCUSSION

We present a large-scale and comprehensive integrative genomic analysis of both localized prostate cancer and mCRPC, with a focus on how structural variation drives each of these clinically distinct disease states. The size of our cohort as well as our harmonized analysis pipeline enable a sharper view of the genetic alterations that drive prostate cancer across its natural history as compared with prior studies, which have involved either smaller cohorts or been limited to a single disease state (9, 13, 15, 59).

In contrast to somatic SNVs/indels and CNAs that occur within coding regions, the functional and clinical significance of alterations within noncoding regions has often been more challenging to interpret, as localized variations in mutability may result in the nomination of certain recurrently mutated sites that do not necessarily drive cancer (11, 12, 44). This issue is even more complex for SVs, in which different classes of SVs spanning the same loci would be predicted to have distinct functional consequences. Our study addresses the former issue by identifying genomic hotspots of structural variation with rigorous correction for covariates including nucleotide composition, replication timing, sensitivity to DNA breaks, repetitive elements, and chromatin state. We address the latter issue by careful curation of SV classes to distinguish those that are likely to be activating versus inactivating (**Figures 1B and 3; Methods**).

Our approach has produced several insights into the recurrent rearrangements that drive prostate cancer. First, several top hotpots of rearrangement genome-wide lie in noncoding regions outside the boundaries of known prostate cancer genes, as previously reported (10, 13, 15). In many cases, such as for *AR*, *MYC*, and *FOXA1*, these hotspots overlap with active chromatin marks and likely represent distal regulatory regions for neighboring prostate cancer genes, as shown by our analyses overlapping SVs with ChIP-Seq on mCRPC specimens (46) (**Figure 2**). These data are intriguing in light of the observation that a majority of prostate cancer germline susceptibility loci are in noncoding regions (60). Second, the loci altered by rearrangements differ across prostate cancer disease states (**Figure 2B**). For example, *TMPRSS2-ERG* rearrangements are enriched in localized prostate cancer versus mCRPC, while alterations in *AR, FOXA1, MYC*, and *LSAMP* are more frequent in mCRPC than in localized disease. Third, certain driver genes are enriched for alteration by SVs as compared to other mutagenic processes. For example, *PTEN* inactivation frequently occurs via gene transecting SV events, while *TP53* inactivation is primarily caused by SNVs (**Figure 1C** **and Figure S1**).

Our systematic genomic discovery efforts confirm the primacy of *AR* as a target of somatic alteration in hormone-refractory mCRPC (13, 15, 26, 53, 61). We have precisely catalogued the diverse genomic mechanisms leading to *AR* activation across our large aggregate cohort and find that different alteration mechanisms are associated with differing levels of *AR* amplification. Whether the precise mechanism by which *AR* is altered in a given patient is associated with differences in response to *AR* pathway inhibition warrants further investigation in clinically annotated cohorts. High levels of *AR* signaling in mCRPC may also underlie the patterns of structural variation seen in this disease state. Strikingly, we found that *AR* binding sites overlapped several of the top SV hotspots in mCRPC (**Figure 2D**), consistent with the notion that androgen signaling may induce DNA double-strand breaks that resolve as rearrangements (62).

In addition to alterations in highly validated prostate cancer genes, we identified highly recurrent rearrangements near or involving genes that have not been extensively studied in prostate cancer in multiple cohorts, such as *LSAMP, PTPRD*, and *TTC28*. *LSAMP* encodes a cell-surface glycoprotein and has a possible tumor suppressor role in several cancers (40–42); notably, deletions near the *LSAMP* locus have been shown in one report to be enriched in African American men with prostate cancer (63). *PTPRD*, a receptor protein tyrosine kinase, has been previously identified as a target of inactivating alteration in glioblastoma (43). We observed frequent SVs near the *TTC28* locus, which encodes an L1 retrotransposon element, specifically in mCRPC (**Figure 1C**). L1 retrotranspositions originating from *TTC28* have been reported previously in colorectal cancer (37–39); our results raise the intriguing possibility that they may also be frequent in prostate cancer, and may be activated by the pressure of hormonal therapy. Interestingly, we also observed SRBs near *ELK4* along with a relatively high frequency of *SLC45A3-ELK4* chimeric transcripts, although it was not clear how the rearrangements at this locus produced the chimeric transcripts in most cases. Whether this fusion functions similarly to or in a distinct mode from other ETS fusions is an exciting area for future study.

Our study also extends beyond the analysis of SVs at individual loci to molecularly subclassify prostate cancers based on their genome-wide signatures of structural variation. Sample clustering based on SV signature exposure defines distinct molecular subtypes of prostate cancer and may find utility alongside signatures of single base substitution and copy number to more precisely define tumor subtypes (55, 64–67). In the mCRPC cohort, we identified 9 molecular subtypes based on SV signature, and several clusters had clear associated genomic alterations including chromoplexy (cluster 1), *BRCA2* alterations (cluster 3), *SPOP* alterations (cluster 5), *TP53* alterations (cluster 6) and *CDK12*/*CCND1* alterations (cluster 7). Future studies with larger WGS and RNA-Seq cohorts may identify associated alterations or transcriptional signatures in the remaining clusters. Notably, unsupervised clustering identified samples with clear SV signatures but without detectable associated mutations in genes or pathways that plausibly contribute to the genomic alterations (**Figure 4**). Moreover, clinical outcomes were more separated by SV signature cluster than by alterations of the mutations associated with those clusters (**Figure S4D-E**).

In sum, these results highlight the dynamic complexity of rearrangements in prostate cancer across disease states and provide insights into new mechanisms of oncogenesis that can be functionally prioritized in future studies. More broadly, our work underscores the key role of large-scale WGS studies in the derivation of a comprehensive molecular taxonomy of prostate cancer.

## METHODS

### Data and code availability

- Whole genome sequencing data have been deposited at dbGaP under accession number phs001577 and access is available upon request.
- All original code has been deposited at GitHub and is publicly available as of the date of publication. Links are provided in the key resources table.
- Any additional information required to reanalyze the data reported in this paper is available from the lead contact upon request.

### Sequence data processing for linked-read genome sequencing data

Data processing of the linked-read genome sequencing data include high molecular weight DNA preparation and sequencing library construction followed protocols as previously described (15). DNA was extracted from tumor samples using the MagAttract HMW DNA Kit (QIAGEN), and then quantified using Quant-it Picogreen assay kit (Thermo Fisher) on a Varioskan Flash Microplate Reader (Thermo Fisher). For germline samples, pre-extracted DNA was size-selected on the PippinHT platform (Sage Science) and then quantified using the Quant-it Picogreen assay kit (Thermo Fisher) on a Varioskan Flash Microplate Reader (Thermo Fisher). Libraries were constructed using the 10X Chromium protocol (10X Genomics), with the fragment sizes determined using the DNA 1000 Kit and 2100 BioAnalyzer (Agilent Technologies) and quantified using qPCR (KAPA Library Quantification Kit, Kapa Biosystems). WGS libraries were sequenced using the Illumina HiSeqX platform. The Long Ranger v2.2.2 pipeline (10X Genomics) was used for aligning sequence reads to the human genome hg38 (GRCh38).

Samples were excluded from the analysis based on having tumor purity less than 15% estimated by TitanCNA or based on cross-individual contamination indicated by SNP fingerprinting. A total of 17 samples with linked-read data was excluded (**Table S1J**).

### List of known prostate cancer driver genes

For analyses limited to established prostate cancer driver genes, a curated list of 159 known prostate cancer driver genes was assembled from several prior studies (10, 13, 15, 24). The list of genes are provided in **Table S1**.

### Somatic mutation analysis

#### Somatic mutation detection

Somatic mutation calls for samples based on linked-read sequencing were generated by Mutect2 from the Genome Analysis Toolkit (GATK) (68). Default parameters were used on individual pairs of tumor and normal samples following the standard GATK pipeline. A panel of normals based on all normal samples was used to filter out germline variants. The SNV calls were further processed using the modified version of LoLoPicker (69) as described previously (15). The panel of normals for LoLoPicker was generated from 52 normal samples based on linked-read sequencing. The final SNV call set was composed of the common variants called by both Mutect2 and LoLoPicker. Somatic indels for linked-read samples were called by Strelka (70). All parameters were default except the following modifications: sindelNoise = 0.000001, minTier1Mapq = 20. Somatic mutation calls for the 101 WGS samples based on short-read sequencing including SNV and indels based on Strelka were obtained from a prior study (13). All variants were further annotated using annovar with “table_annovar.pl” to functionally annotate genetic variants. The parameter - neargene was set to 5000 to define the promoter region as 5 kB upstream of the transcription start site of a protein coding gene.

#### Analysis of significantly mutated genes

R package dndscv (71) was used to identify significantly mutated genes. For driver discovery on GRCh38, a precomputed database corresponding to human genome GRCh38.p12 was downloaded and used as the reference database. A global q-value ≤ 0.1 was applied to identify statistically significant (novel) driver genes. To reduce false positives and increase the signal to noise ratio, we only considered mutations in Cancer Gene Census genes (v81) (72).

### Copy-number analysis of linked-read WGS and short-read WGS data

#### Copy-number calls

The ploidy and purity corrected copy-number of all mCRPC samples in this study was analyzed by TitanCNA (73) and ichorCNA (74), with different pipeline settings. For WCDT samples, the snakemake workflow for Illumina sequencing was applied with the following parameters modified: ichorCNA_normal: c(0.25, 0.5, 0.75); ichorCNA_ploidy: c(2,3,4); ichorCNA_includeHOMD: TRUE; ichorCNA_minMapScore: 0.75; ichorCNA_maxFracGenomeSubclone: 0.5; ichorCNA_maxFracCNASubclone: 0.7; TitanCNA_maxNumClonalClusters: 3; TitanCNA_maxPloidy: 4. The workflow is available at https://github.com/GavinHaLab/TitanCNA_SV_WGS.

For linked-read data samples, a Snakemake workflow for 10X Genomics whole genome sequencing data was used with the following parameters modified: TitanCNA_maxNumClonalClusters: 3; TitanCNA_maxPloidy: 4. TitanCNA solutions were generated for number of clonal clusters from 1 to 3 and ploidy initializations from 2 to 4. Optimal solutions were selected as described, with manual inspection to confirm tumor ploidy and clonal cluster selection (15); solutions are provided in **Table S1J**. The workflow can be accessed at https://github.com/GavinHaLab/TitanCNA_10X_snakemake. The final copy-number call-set is included in **Table S1I**.

#### Recurrent somatic copy-number alteration

GISTIC 2.0 was used to detect regions with recurrent CNA in mCRPC samples. For input, all copy numbers (logR_Copy_Number from TITAN output) were converted to log2 copy ratio using the median logR copy number from genome-wide (separately for autosomes and X chromosome) as denominator. We set corrected logR copy number to -1.5 for segments where corrected log R copy number below -1.5 and set values to 0 if copy neutral. GISTIC2.0 was run with the following parameters: td 0.5; ta 0.1; genegistic 0; maxseg 5000; js 4; cap 1.5; broad 1; brlen 0.75; conf 0.99; qvt 0.25; armpeel 1; rx 0; gcm mean; do_gene_gistic 1; savegene 1; scent median. Wide peaks detected by GISTIC2 were re-annotated based on overlapping genomic coordinates, using prostate cancer driver genes.

### Structural variant analysis

#### Structural variant detection in linked-read and short-read whole genome sequencing data

For each tumor-normal pair of samples with linked-read genome sequencing data, three variant callers were used to detect structural variants: SvABA (75), GROC-SVS (76), Long Ranger version 2.2.2 (https://support.10xgenomics.com/genome-exome/software/pipelines/latest/using/wgs).

The SvABA analysis was performed using default tumor-normal paired settings. Re-analysis of low confidence (based on evidence from discordant and split reads) events filtered by SvABA was performed to ‘rescue’ SVs using linked-read barcode overlap between pairs of breakpoints within a given SV event, as previously described (15). Only SV events having span of 1.5 times the mean molecule length in the library were considered for rescue. We further rescued low confidence intra-chromosomal SV events with span > 50 kB filtered by SvABA if at least one of the breakpoint pair was within 100 kB of a CNA boundary or (2) if both breakpoints were each within 1 Mb of the boundaries for the overlapping CNA event and the length of the SV overlaps this CNA event by > 75%. Inter-chromosomal translocation SV events filtered by SvABA are rescued if both breakpoints were within 100 kB of CNA boundaries.

GROC-SVS analysis was performed using two-sample (tumor-normal paired) mode or three-sample (pre-treatment, post-progression, normal) mode when applicable. SV events were retained if all following conditions were satisfied: (1) p < 1x10^-10^, (2) minimum barcode overlap ≥ 2 on the same haplotype, (3) no more than 1 barcode overlap between different haplotypes, (4) FILTER value reported by the software was within this set {“PASS”, “NOLONGFRAGS”, “NEARBYSNVS”, or “NEARBYSNVS; NOLONGFRAGS”}, and (5) classified as somatic.

Long Ranger analysis generated SV calls for tumor and normal samples, independently. For each tumor-normal pair, both large SVs (“large_sv_calls.bedpe”) and deletions (“dels.vcf”) were combined for individual samples. Somatic tumor SVs were determined as events that were not found in the matched normal sample based on the left breakpoints in tumor and normal being within 1 kB and the right breakpoints in tumor and normal samples being within 1 kB. Only SV events with FILTER values within this set {“PASS”, “LOCAL_ASM”, “SV”, “CNV, SV”} and intra-chromosomal events with span ≥ 100 kB were considered. SV events were only retained if both breakpoints of an SV event were within 500 kB the boundaries of an overlapping CNA event and the length of SV overlaps this CNA event by > 75%.

SV events from these three callers were then combined by taking the union of the filtered events from. Intersecting events between 2 or more call-sets were determined if both breakpoints of one event were located within 5 kB from both breakpoints of the event detected by the other tool. Then the details of this event were retained based the priority ordered by SvABA, GROC-SVS, Long Ranger. Long Ranger SV events were further filtered out if they were not intersecting events detected by at least one other tool. SV events with span less than 1 kB were excluded from downstream analyses.

An SV panel of normals (PoN) was generated using germline events from SvABA and Long Ranger calls. There are two components to this panel: (1) frequency of germline events at exact breakpoint locations (SVpon.bkpt) and (2) frequency of germline event breakpoint overlapping within tiled windows of 1 kB (SVpon.blackListBins). The PoN was used to filter events in the combined SV call-set when an SV has at least one breakpoint with SVpon.bkpt ≥ 2 and overlapping bin with SVpon.blackListBins ≥ 100.

The workflow for SV analysis from linked-read sequencing data can be accessed at https://github.com/GavinHaLab/SV_10X_analysis. Manual curation of filtered SV events in the *AR* locus was performed and rescued events were labeled “Manual”. The final SV call-set is included in **Table S1K**.

For samples based on short-read WGS, SvABA was used in tumor-normal paired mode for SV detection with default parameters. Intra-chromosomal SV events with span > 1 kB were retained. The SvABA workflow can be accessed at https://github.com/GavinHaLab/TitanCNA_SV_WGS

#### Classification of structural variants in mCRPC

SV types were annotated based on orientations of breakpoints and bin-level copy-number around breakpoints. The orientation of one breakpoint was defined based on the fragment of DNA molecule being connected to the altered molecule. If the connected fragment was to the 5’-end of the breakpoint, *i.e.*, “upstream” or “left” to the breakpoint, then the orientation was annotated as forward or “+”; on the contrary, if the connected fragment was located to the 3’-end of the breakpoint, the orientation was annotated as reverse or “-”. The copy-number near each breakpoint was evaluated using 10 kB bins. For one SV event, copy-number values of the bins located to the upstream and downstream of breakpoint 1 were denoted as *c_1_^up^* and *c_1_^down^*, respectively; similarly, the copy-number values for breakpoint 2 were denoted as *c_2_^up^* and *c_2_^down^*. In addition, then mean copy-number *c^mean^* of the 10 kB bins between the two breakpoints of one SV event and the number of bins *s* were also considered during SV classification. Intra-chromosomal SV events, *i.e.*, both breakpoints were located on the same chromosome, were classified to the list of SV types below following the corresponding classification criteria.

- Deletion. Events having the orientation combination (reverse, forward) and length between 10 kB and 1 Mb were classified as deletions. The copy-number values of breakpoints should satisfy *c_1_^up^* > *c_1_^down^* or *c_2_^up^* < *c_2_^down^*, and *c_1_^up^* > *c^mean^* or *c_2_^down^* > *c^mean^*, and *s* ≤ 5. In addition, events overlapping copy-number deletion or LOH segments were also considered as deletions.
- Tandem duplication. Events having the orientation combination (forward, reverse) and length between 10 kB and 1 Mb were classified as tandem duplications. The copy-number values of breakpoints should satisfy *c_1_^up^* < *c_1_^down^* or *c_2_^up^* > *c_2_^down^*, and *c_1_^up^* < *c^mean^* or *c_2_^down^* < *c^mean^*, and *s* ≤ 5. In addition, events overlapping copy-number gain or copy neutral LOH segments were also considered as tandem duplications.
- Inversion. Events having the orientation combination (forward, forward) or (reverse, reverse) and length between 10 kB and 5 Mb were classified as inversions. Furthermore, inversion events shorter than 30 kB with unequal copy-numbers around either breakpoint were classified as fold-back inversions.
- Balanced rearrangement (balanced). Events having the orientation combination same to inversion (forward, forward) or (reverse, reverse), but length larger than 5 Mb were classified as balanced events. The copy-number values of breakpoints should satisfy *c_1_^up^* = *c_1_^down^* and *c_2_^up^* = *c_2_^down^*, or *c_1_^up^* = *c^mean^* and *c_2_^down^* = *c^mean^*.
- Unbalanced rearrangement (unbalanced). Intra-chromosomal events which did not fulfill any of the above criteria and having length larger than 10 kB were classified as unbalanced events.
- All SV events with two breakpoints located on different chromosomes were classified as translocations.

#### ICGC/TCGA PCAWG localized prostate cancer structural variants

We obtained localized prostate cancer structural variant calls from ICGC Data Portal release 28 (https://dcc.icgc.org/releases/PCAWG/consensus_sv). In this consensus SV file, each SV event was predicted by at least two variant callers. Samples that were classified as prostate adenocarcinoma (PRAD) and early onset prostate cancer (EOPC) were selected. A total of 278 samples successfully lifted over to genome build GRCh38. To maximize consistency with mCRPC datasets, we used only the PCAWG consensus SVs that included “SNOWMAN” as one of the tools. Note that “SNOWMAN” was the previous name for SvABA. Intrachromosomal SV events shorter than 10 kB were excluded.

We obtained ICGC early-onset PC (EOPC) and late-onset (LOPC) structural variant calls from Gerhauser et al, 2018 (18) (https://data.mendeley.com/datasets/6gtrrxrn2c/1). This dataset includes a total of 253 samples, consisting of 206 EOPC and 47 LOPC. We used the SV calls generated in the original study using DELLY2 (Rausch et al., 2012) PCAWG analysis workflow (https://github.com/ICGC-TCGA-PanCancer/pcawg_delly_workflow). SV coordinates were lifted over to GRCh38.

### Tandem duplicator phenotype

For all samples in the combined cohort, the TDP status was predicted using copy-number and SV by counting the number of copy-number segments overlapping with tandem duplication SV events, *i.e.,* gain segments. A sample was considered as TDP if it has more than 300, or 90 gain segments for samples based on linked-read sequencing and short-read sequencing, respectively. The number of segments with gain and median length SV are reported in **Table S1L**.

### Chromothripsis analysis

Chromothripsis events were detected by ShatterSeek R package (77). Structural variants calls by SvABA and copy-number calls by TitanCNA were used as input data (excluding Y chromosome). In the input, consecutive segments were joined as one if they had the same copy-number value and centromere regions were filtered out.

Manual inspection was performed for reported chromothripsis-like events after adapting criteria thresholds. For samples based on short-read sequencing, confidence classification criteria were refined from the ShatterSeek documentation. Following criteria were used for high confidence calls: total number of intra-chromosomal structural variants events involved in the event ≥ 10; max number of oscillating CN segments (two states) ≥ 10; satisfying either the chromosomal enrichment or the exponential distribution of breakpoints test (p ≤ 0.05). For samples based on linked-read sequencing, we filtered these calls based on a weighted score that is primarily determined by the number of SVs in a cluster, with less weight given to CN oscillations. In this analysis, events with a score over 0.8 were considered as high confidence and all other events were excluded. The score is defined based on the following terms (ranges from 0 to 1).

- Weight 0.6 if total number of intra-chromosomal structural variants events involved in the event ≥ 10.
- Weight 0.2 for max number of oscillating CN segments (two states) ≥ 7 or max number of oscillating CN segments (three states) ≥ 14.
- Weight 0.1 for passing chromosomal enrichment test by ShatterSeek.
- Weight 0.1 for passing exponential distribution of breakpoints test.

### Chromoplexy analysis

ChainFinder was used to detect chromoplexy events (8). Ten samples that were considered as TDP (01115374-TA2, 01115202-TC2, 01115248-TA3, 01115503-TC2, 01115257-TA4, 01115284-TA9, 01115414-TA1, DTB-063-BL, DTB-183-BL, DTB-214-BL) were excluded from this analysis. In addition, four samples that were found to cause numeric instabilities of ChainFinder were also excluded (DTB-023-BL, DTB-102-PRO, DTB-111-PRO, DTB-151-BL). The SV calls of remaining samples were further filtered to exclude those that were located within 5 Mb from chromosomal ends or overlapping chromothripsis regions. For copy-number input, segments that were determined as copy neutral by TitanCNA were set to have log copy-ratio of 0. Copy-ratio of the other segments were computed from copy-number values generated by TitanCNA divided by 2 for autosomes or 1 for X chromosome. Log copy-ratio values less than - 1.5 were set to -1.5. The output of ChainFinder was used for determining chromoplexy status of individual samples. A chromoplexy event was defined as a chain including at least 5 rearrangement events and involving more than 2 different chromosomes. Samples having at least 2 such events were considered positive for chromoplexy status.

### ChIP-seq data analysis

ChIP-seq data used in this study were downloaded from Gene Expression Omnibus (GEO) (78, 79) and the Sequence Read Archive (SRA) (1). Short reads were mapped to the human genome GRCh38 (hg38) using bwa (80). Because read lengths were less than 50bp, the bwa aln command with default parameters was used for mapping. MACS2 (81) was used to identify peaks from mapped ChIP-seq data. For histone modification marks, MACS2 callpeak command was applied with --nomodel --broad –extsize 146. For CTCF data, MACS2 callpeak command was used with --nomodel --extsize 200. Below is the list of ChIP-seq datasets involved in this analysis.

- H3K4me3, H3K27me3 and CTCF (GSE38685) (82).
- H3K36me3 and H3K9me3 (GSE98732) (83).
- H3K4me1 and H3K27ac (GSE73785) (84).

For *AR* binding site (ARBS), the peak files were downloaded from two different datasets and converted to hg38 coordinates. For primary prostate cancer, ARBS data were downloaded from GSE70079 (Pomerantz et al., 2015). The union of all tumor sample peaks was used. For mCRPC, met-specfic ARBS data were obtained from a previous study (46).

### Identification of SRB regions

#### Masking the human genome based on mappability

The human genome was divided into 100 kB non-overlapping bins for detection of significantly recurrent breakpoint regions (SRB). A low-mappability mask was generated for the hg38 genome to screen out out regions that are difficult for variant calling based on short-read sequencing. We adopted procedures from a previous study (86) to construct a mask corresponding to regions with low mappability in the human genome. Below is a list of masked regions included in the low-mappability mask.

- Composition mask. This set of masked regions includes regions with low sequence complexity detected by mdust, regions with long homopolymers detected by seqtk, satellite regions annotated by RepeatMasker (87), and low complexity regions annotated by RepeatMasker.
- Mappability mask. This mask was based on mappability of *k*-mers in the human genome hg38. The value *k* was set to 75 which is half of the read length of WGS data in this study. Each base in the genome was assigned a mappability level, based on the mapping ambiguity of all 75-mers overlapping this specific base. See below for the list of mappability levels. Regions with mappability level 0 and 1 were included in the low-mappability mask.

- Level 0: all 75-mers overlapping this base could not be mapped to the genome uniquely.
- Level 1: more than 50% of overlapping 75-mers are not uniquely mapped.
- Level 2: more than 50% of overlapping 75-mers are uniquely mapped with 1-mismatch hits.
- Level 3: more than 50% of overlapping 75-mers are uniquely mapped without 1-mismatch hits.

In addition, we used GATK CallableLoci to mark regions with high confidence of variant detection based on coverage. Together, the intersection of unmasked regions and callable loci were defined as the eligible territories for SRB detection. The 100 kB bins with less than 75% overlap with eligible territories were excluded from the analysis.

#### Generating covariates for regression analysis

To accurately model the genomic features of mCRPC, we incorporated the following covariates.

- Nucleotide composition, including GC content, CpG fraction and TpC fraction per 10 kB non-overlapping bin in the genome.
- Replication timing of LNCaP (data obtained from ENCODE under accession ENCFF995YGM, lifted over from hg19 to hg38) (88, 89).
- DNase I hypersensitive sites (data obtained from ENCODE under accession ENCFF434GSJ, lifted over to hg38).
- Repeats annotated by RepeatMasker, including LINE, SINE, LTR, DNA transposon and simple repeats.
- Heterochromatin regions inferred by ChromHMM (90) with the 18-state model parameters from the Roadmap Epigenomics Project (91), based LNCaP ChIP-seq data of H3K4me1, H3K4me3, H3K4ac H3K27me3, H3K36me3 and H3K9me3.
- Common fragile sites downloaded from HGNC biomart (92).

#### SRB detection

Structural variants from the final call set were used for statistical enrichment of recurrent breakpoints within 100 kB bins using a Gamma-Poisson regression implemented in the package, fish.hook (44). Breakpoints of SVs were treated independently. The Benjamini-Hochberg procedure was used for multiple testing correction and bins with q-value ≤ 0.1 were determined to be significant. The distances of individual known driver genes to those significant bins were evaluated based on the shortest genomic distance between the gene and bin boundaries, regardless of gene orientations.

### Annotation of gene alteration status

#### Gene alteration by copy-number

Copy-number segments were excluded if their cellular fraction was lower than 0.8, except for those which were determined as copy neutral or copy-number greater than 4. The gene annotation was based on known protein coding genes from GenCode release 30 (GRCh38.p12) (93). For each gene, its copy-number was assigned to the copy-number value and LOH status of the segment that has the largest overlap with it. The gene-level copy-number was normalized based on ploidy of the corresponding sample, with autosomal genes normalized by the inferred ploidy rounded to nearest integer, and X-linked genes normalized by half such value. Then the copy-number status of each gene was categorized based on the following criteria.

- Amplification. Normalized gene-level copy-number is greater than or equal to 2.5.
- Gain. Normalized gene-level copy-number is between 2 and 2.5.
- Homozygous deletion. Normalized gene-level copy-number is 0.
- Deletion with LOH. Normalized gene-level copy-number is between 0 and 1, and LOH status was found.
- Copy neutral LOH. Normalized gene-level copy-number is 1 and LOH status was found.

#### Gene alteration by structural variant

Gene coordinates were based on ENSEMBL v33 of hg38 (94). Gene body region of one gene was defined as the widest region of all known isoforms collapsed. Gene flanking region was defined as the corresponding two 1 Mb regions next to the gene body region on 5’-end and 3’-end, respectively.

Gene alteration status by genome rearrangements was defined based on the breakpoints and directions of involving structural variant events. A gene in one WGS sample (gene-sample pair) was considered having gene transecting events if any breakpoints of SV events were located within the gene body region. If the gene transecting status did not apply, then this gene-sample pair was examined for gene flanking status if the breakpoints of any intra-chromosomal SV events, including tandem duplications, deletions, and inversions, were located within the gene flanking regions. Additionally, translocation events including intra-chromosomal balanced and unbalanced events which spanned over 10 Mb, and inter-chromosomal translocation events were considered altering the gene flanking regions if any of their breakpoints was in the gene flanking region, and the direction of the SV was going towards the gene body region. The alteration status of rearrangements for each gene-sample pair was exclusive between gene transecting and gene flanking, with the former being prioritized in report.

#### AR alteration analysis

Copy-number of the *AR* gene (chrX:67,544,623-67,730,619) and the *AR* enhancer region (chrX:66,895,000-66,910,000) were each computed as the mean corrected total copy-number across the 10 kB bins overlapping each region. The copy-number was further normalized by sample ploidy as previously described. Amplification status of *AR* was determined by comparing the log2 fold-change *FC* of enhancer-level over gene-level copy-number. Four distinct groups were defined based on copy-number and *FC* as below.

- Co-amplification. Ploidy normalized copy-number values of both *AR* gene body and enhancer are greater than 1.5.
- Selective *AR* amplification. *FC* < -log2(1.5) and enhancer copy-number is less than 1.5.
- Selective enhancer copy gain. *FC* > log2(1.5) and *AR* gene body copy-number is less than 1.5.
- Lack of amplification for both. All other cases were considered as no amplification for both regions.

ANCOVA test was used to test if different patterns of *AR* amplification have an impact on *AR* expression. See Statistics section.

### Gene expression

TPM values for a subset of the samples based on linked-read sequencing were obtained from cBioportal (95, 96). For samples based on short-read sequencing the TPM values were obtained from a previous study (13).

### Gene fusion analysis

Fusion status of the main members of the ETS family, including *ERG*, *ETV1*, *ETV4*, *ETV5* and *ELK4* was analyzed. Determination of gene fusion status was based on both DNA and RNA levels. For DNA, structural variants transecting gene body regions were used. SV events were considered supporting gene fusion only if they satisfy the following criteria: (1) the breakpoints of this event must be located within the ETS gene and another protein coding gene, respectively; (2) the orientation of the breakpoint located within the ETS gene must be pointing towards the coding sequence of ETS domain. For RNA, arriba was used to detect fusion transcripts from RNA-seq data (97). The fusion status was only confirmed if all following conditions were satisfied: (1) the complete ETS domain was included in the fusion product; (2) detection confidence reported by arriba is “high”; (3) coding sequence in the fusion transcript was in sense orientation and no out-of-frame shifts.

### SV signature analysis

#### Signature extraction and clustering

*De novo* signature extraction was performed on all SV events called by SvABA of the combined cohort using signature.tools.lib (55) with the recommended settings of 20 bootstraps, 200 repeats, the clustering with matching algorithm, the KLD objective function, and RTOL = 0.001. The exposure of one signature in one sample is defined as the median activity of the signature within the sample across all bootstraps. For clustering, the reference signature exposure values for each sample based on short-read sequencing were normalized such that the sum of exposure values per sample is 1, and the normalized exposure values for each signature were mean-centered across all samples. A Euclidean distance matrix was computed and then samples were clustered with the Ward.D2 algorithm using R’s hclust function. We chose the number of clusters to be k = 9 based on dendrogram using cutree function in R.

### STATISTICS

#### Association of AR locus amplification status and AR expression

ANCOVA test was used to test if different patterns of *AR* amplification have an impact on *AR* expression. Batch corrected log10(TPM+1) values using ComBat from sva R package (v3.34.0) were used for *AR* expression level. We fit the ANCOVA model using *AR* expression as the response variable, *AR* amplification status as the predictor variable, and ploidy, purity as covariates. The function Anova in the car package (v3.0-5) was used with Type III sum of squares for the model. Post hoc analysis was performed to determine the specific differences among four different *AR* amplification status. The function glht was used within the multcomp package (v1.4-11) in R to perform Tukey’s Test for multiple comparisons.

#### Enrichment of alterations in SV clusters

All 9 identified SV clusters were analyzed for enrichment of alterations. To make the analysis unbiased by SV signature, we limited our search to alteration types that were orthogonal to rearrangements, which include SNV, copy-number gain and copy-number loss. We performed hypothesis testing on each driver-alteration pair, and also on chromoplexy and chromothripsis. For each SV cluster, a χ^2^ test was performed for each driver gene alteration status, with samples within group being tested against samples belonging to all 8 other SV clusters. Multiple testing adjustment based on Benjamini-Hochberg FDR was performed to compute q-values. Alteration categories with q-values less than 0.25 were determined as enriched in the corresponding SV cluster.

#### Survival analysis

Survival data was obtained from (98). Survival analyses were conducted using the Kaplan-Meier method with log-rank testing for significance. The function survfit from survival R package was used to perform the analysis.

## STUDY APPROVAL

For tumor biopsies profiled via linked-read sequencing, samples were collected from individuals with mCRPC who provided informed consent on institutional IRB-reviewed protocols, as previously described (15). Uniformly reanalyzed data were generated as described in the respective studies (9, 13, 18).

## AUTHOR CONTRIBUTIONS

**Conceptualization:** M-E.T, M.M., S.R.V., G.H.

**Methodology:** M.Z., M.K., S.R.V., G.H.

**Software:** M.Z., M.K., A.C.H., G.H.

**Formal Analysis:** M.Z., M.K., A.C.H., K.L., Y.L., M.R., W.H., J.C-Z, S.R.V., G.H.

**Data Curation:** M.Z., M.K., A.C.H., Z.Z., S.R.V., G.H.

**Writing – Original Draft:** M.Z., M.M., G.H., S.R.V.

**Writing – Review & Editing:** M.Z., R.B., E.M.V., A.D.C. P.S.N., M.L.F., M-E.T., M.M., G.H., S.R.V. **Visualization:** M.Z., M.K., A.C.H., S.R.V., G.H.

**Supervision:** M-E.T., M.M., S.R.V., G.H.

**Funding Acquisition:** S.R.V., G.H., M.M.

## ACKNOWLEDGEMENTS

We thank the many patients and their families for their generosity in contributing to this study. We also thank the Prostate Cancer Foundation (PCF) and Stand Up 2 Cancer (SU2C) International Prostate Cancer Dream Team for contributions to specimen acquisition.

This work was supported by the National Institutes of Health (K22 CA237746 to G.H.; P01 CA163227 and R01 CA234715 to P.S.N.; R01 GM107427, R01 CA193910, and R01 CA251555 to M.L.F.; R35 CA197568 to M.M.), Department of Defense Prostate Cancer Research Program (Physician Research Award W81XWH-17-1-0358 to S.R.V.; W81XWH-19-1-0565 and W81XWH-21-1-0234 to M.L.F.; PC200262 to P.S.N.), PCF Young Investigator Awards (G.H. and S.R.V.), PCF-Movember Challenge Award (to E.M.V.), Brotman Baty Institute for Precision Medicine (to G.H.), the Fund for Innovation in Cancer Informatics Major Grant (to G.H.), the V Foundation Scholar Grant (to G.H.), Wong Family Award in Translational Oncology and Dana-Farber Cancer Institute Medical Oncology grant (to A.D.C.), H.L. Snyder Medical Research Foundation and the Cutler Family Fund for Prevention and Early Detection (to M.L.F.), and the Pan-Mass Challenge team IMAGINE (to M-E.T.), American Cancer Society Research Professor (M.M.).

This research was also supported in part by the NIH/NCI Cancer Center Support Grant P30 CA015704, Pacific Northwest Prostate Cancer SPORE (P50 CA097186), and Scientific Computing Infrastructure (ORIP Grant S10OD028685).

