## Supplementary Materials for "Patterns of Structural Variation Define Prostate Cancer Across Disease States"

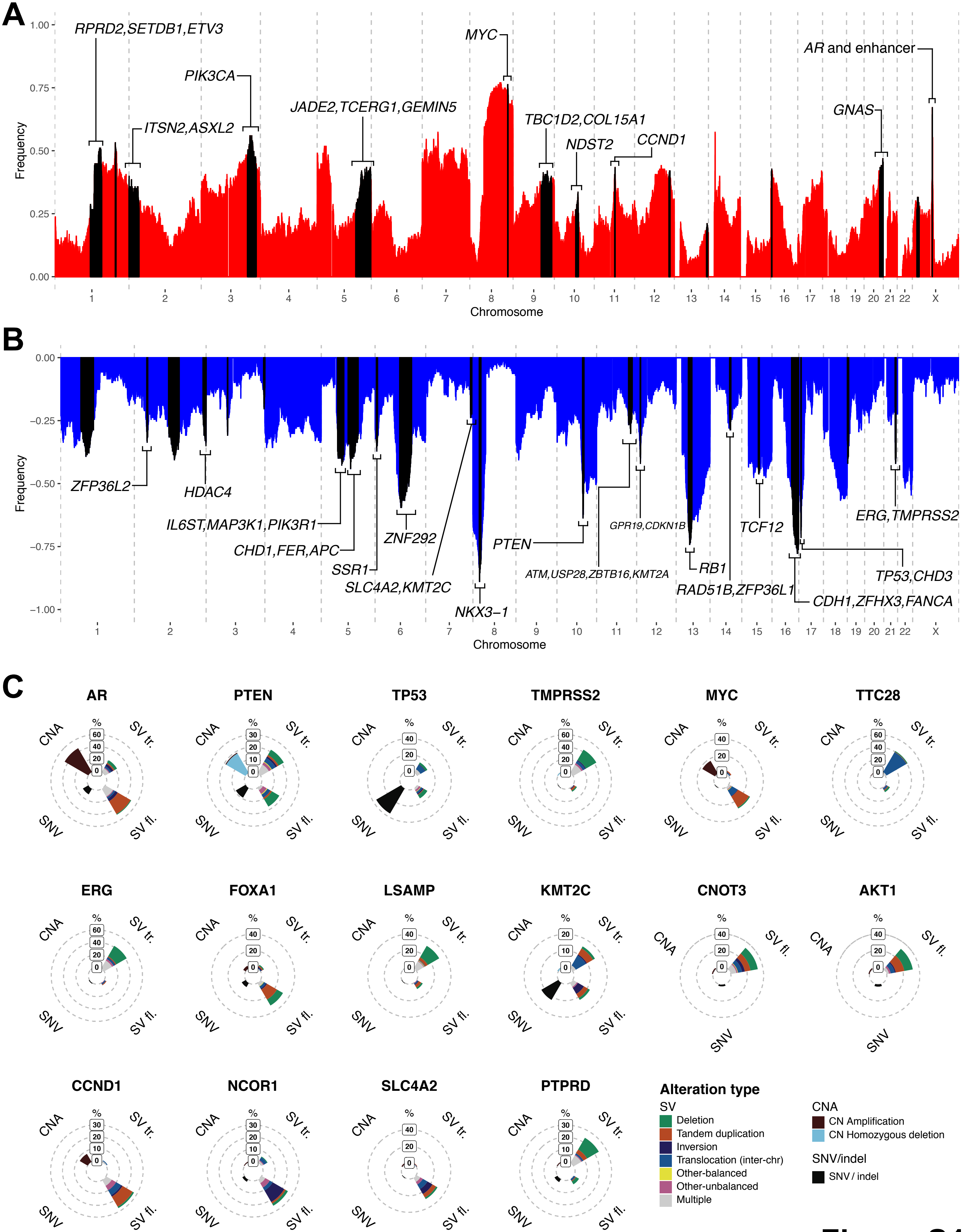
A
RPRD2,SETDB1,ETV3
ITSN2,ASXL2
PIK3CA
JADE2,TCERG1,GEMIN5
MYC
TBC1D2,COL15A1
NDST2
CCND1
GNAS
AR and enhancer
Frequency
Chromosome
B
ZFP36L2
HDAC4
IL6ST,MAP3K1,PIK3R1
CHD1,FER,APC
SSR1
SLC4A2,KMT2C
ZNF292
NKX3−1
PTEN
ATM,USP28,ZBTB16,KMT2A
GPR19,CDKN1B
RB1
RAD51B,ZFP36L1
TCF12
CDH1,ZFHX3,FANCA
TP53,CHD3
ERG,TMPRSS2
Frequency
Chromosome
C
AR
PTEN
TP53
TMPRSS2
MYC
TTC28
ERG
FOXA1
LSAMP
KMT2C
CNOT3
AKT1
CCND1
NCOR1
SLC4A2
PTPRD
CNA
SV tr.
SV fl.
SNV
Alteration type
SV
Deletion
Tandem duplication
Inversion
Translocation (inter-chr)
Other-balanced
Other-unbalanced
Multiple
CNA
CN Amplification
CN Homozygous deletion
SNV/indel
SNV / indel

**Figure S1**

**Figure S1. Recurrent CNA and alteration profiles of most frequently altered genes.**

**(A)** Recurrent copy number gain events in the genome. The frequencies of copy number gain are plotted in red according to their genomic coordinates. Regions with significantly recurring CNA are colored in black. Known driver genes that are within those regions are labeled.

**(B)** Recurrent copy number loss events in the genome. The frequencies of copy number loss are plotted in blue with y-axis inverted.

**(C)** Alteration profiles of known prostate cancer driver genes. Alterations are categorized into CNA, SNV and SV, with SV being further divided into gene transecting (SV tr.) and gene flanking (SV fl.). The percentages of samples carrying corresponding alterations are shown as stacked bars. All known prostate cancer driver genes were considered and the top 16 genes with overall alteration frequencies above 30% are shown.

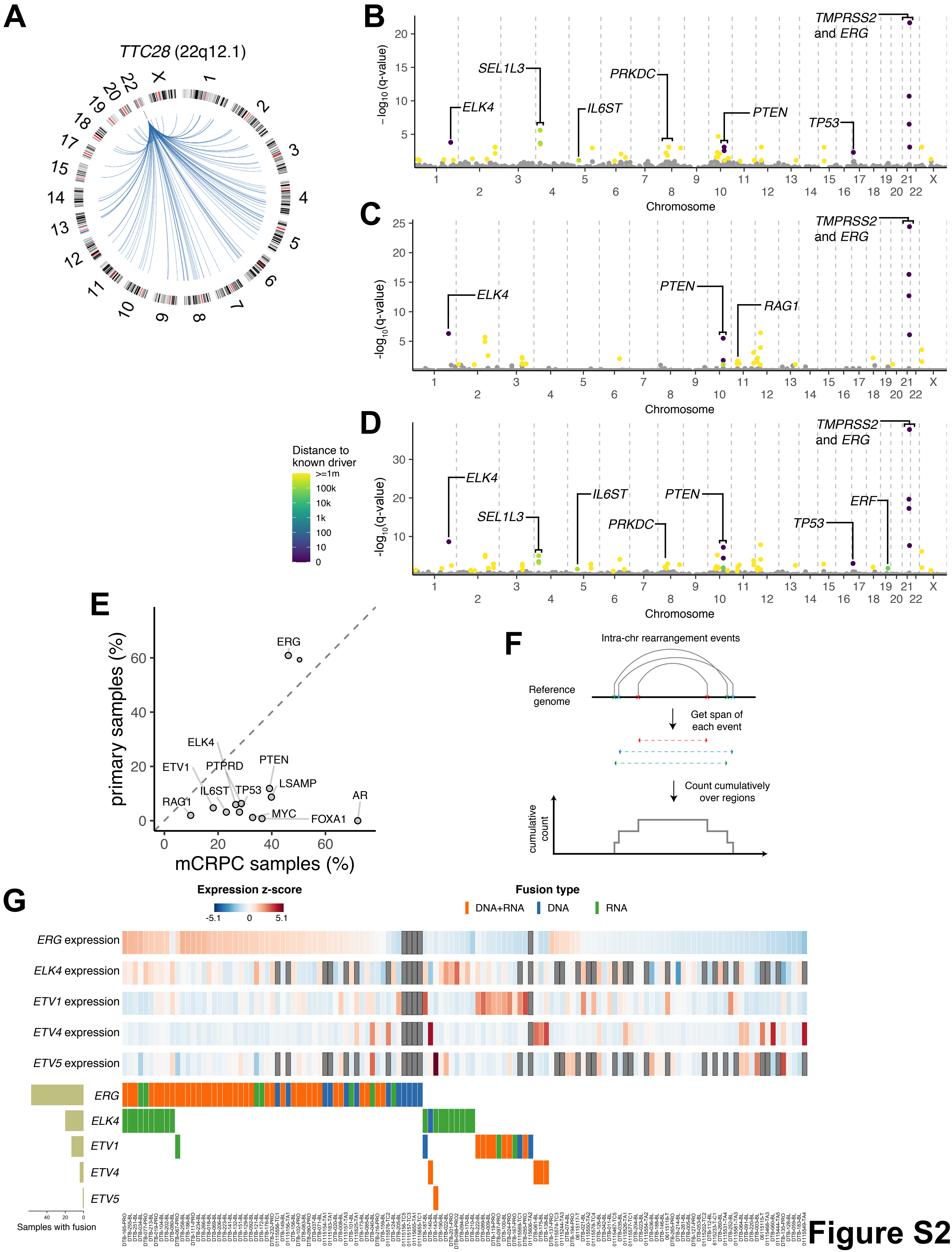
A
TTC28 (22q12.1)
B
TMPRSS2 and ERG
SEL1L3
PRKDC
ELK4
IL6ST
PTEN
TP53
−log10(q-value)
Chromosome
C
TMPRSS2 and ERG
ELK4
PTEN
RAG1
-log10(q-value)
Chromosome
D
TMPRSS2 and ERG
ELK4
IL6ST
PTEN
SEL1L3
PRKDC
ERF
TP53
-log10(q-value)
Chromosome
Distance to known driver
>=1m
100k
10k
1k
100
10
0
E
primary samples (%)
mCRPC samples (%)
ERG
ELK4
ETV1
PTPRD
PTEN
LSAMP
IL6ST
TP53
RAG1
MYC
FOXA1
AR
F
Intra-chr rearrangement events
Reference genome
Get span of each event
Count cumulatively over regions
cumulative count
G
Expression z-score
Fusion type
DNA+RNA
DNA
RNA
ERG expression
ELK4 expression
ETV1 expression
ETV4 expression
ETV5 expression
ERG
ELK4
ETV1
ETV4
ETV5
Samples with fusion

Figure S2

### Figure S2. Recurrent SV in localized prostate cancer and landscape of ETS fusion in mCRPC.

**(A)** Translocation events originating from *TTC28*. In the circos plot, *TTC28* is labeled with a vertical bar at the 22q12.1 locus. Translocation events which have breakpoints located within 5 kB to the 3'-end of the L1 retrotransposon are visualized as blue arcs.

**(B)** SRBs detected in the cohort of localized prostate cancers from the ICGC Pan-Cancer Analysis Working Group (PCAWG) (n=278). The criteria for coloring and labeling are the same as Figure 2.

**(C)** SRBs detected in the cohort of localized prostate cancers from ICGC/Gerhauser dataset. (n=253)

**(D)** SRBs detected in localized prostate cancers from the combined cohorts from PCAWG (n=278) and ICGC/Gerhauser dataset (n=253).

**(E)** Comparison of SV alteration frequency in mCRPC versus primary localized prostate cancer. The union set of genes (n=13) within 1 Mb of SRB hotspot regions in mCRPC and localized prostate cancer (ICGC/Gerhauser) dataset was included in the comparison. The frequencies represent total gene transecting and flanking SV events. All labeled genes were significantly enriched in either mCRPC or primary localized tumors (Fisher's test, $p\text{-value} < 0.05$).

**(F)** Schematics of cumulative counts from intra-chromosomal SV events. Individual SV events are indicated by a grey arc, and colored crosses correspond to breakpoints of each event.

**(G)** Expression and fusion status for main genes of the ETS family. The expression values were normalized from TPM to z-score within each gene. Grey boxes indicate expression data are not available. For fusion status, color indicates the data type which was used to call fusion.

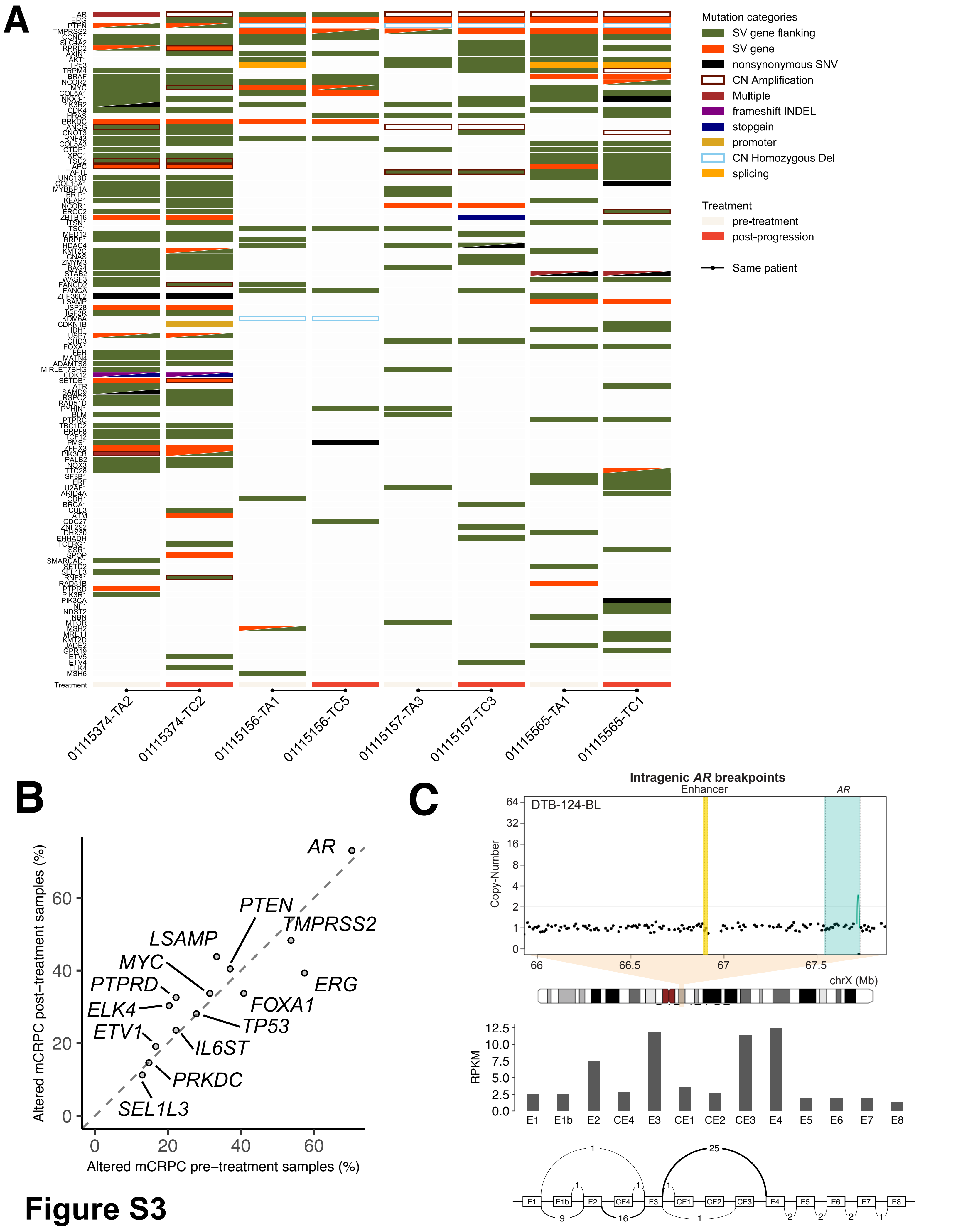
A
AR
ERG
PTEN
TMPRSS2
CCND1
SLC4A2
RPRD2
AXIN1
AKT1
TP53
TRPM4
BRAF
NCOR2
MYC
COL5A1
NKX3-1
PIK3R2
CDK4
HRAS
PRKDC
FANCG
CNOT3
RNF43
COL5A3
CTDP1
XPO1
TSC2
APC
TAF1L
UNC13D
COL15A1
MYBBP1A
BRIP1
KEAP1
NCOR1
ERCC2
ZBTB16
ITSN1
TSC1
MED12
BRPF1
HDAC4
KMT2C
GNAS
ZMYM3
BAG4
STAB2
WASF3
FANCD2
FANCA
ZFP36L2
LSAMP
USP28
IGF2R
KDM6A
CDKN1B
IDH1
USP7
CHD3
FOXA1
FER
MATN4
ADAMTS8
MIRLET7BHG
CDK12
SETDB1
ATR
SAMD9
RSPO2
RAD51D
PYHIN1
BLM
PTPRC
TBC1D2
PRPF8
TCF12
PMS1
ZFHX3
PIK3CB
PALB2
NOX3
TTC28
SF3B1
ERF
U2AF1
ARID4A
CDH1
BRCA1
CUL3
ATM
CDC27
ZNF292
DHX30
EHHADH
TCERG1
SSR1
SPOP
SMARCAD1
SETD2
SEL1L3
RNF31
RAD51B
PTPRD
PIK3R1
PIK3CA
NF1
NDST2
NBN
MTOR
MSH2
MRE11
KMT2D
JADE2
GPR19
ETV5
ETV4
ELK4
MSH6
Treatment
01115374-TA2
01115374-TC2
01115156-TA1
01115156-TC5
01115157-TA3
01115157-TC3
01115565-TA1
01115565-TC1
Mutation categories
SV gene flanking
SV gene
nonsynonymous SNV
CN Amplification
Multiple
frameshift INDEL
stopgain
promoter
CN Homozygous Del
splicing
Treatment
pre-treatment
post-progression
Same patient
B
Altered mCRPC post−treatment samples (%)
Altered mCRPC pre−treatment samples (%)
0
20
40
60
AR
PTEN
TMPRSS2
LSAMP
MYC
PTPRD
ERG
FOXA1
ELK4
TP53
ETV1
IL6ST
PRKDC
SEL1L3
C
Intragenic AR breakpoints
Enhancer
AR
DTB-124-BL
Copy-Number
64
32
16
8
4
2
1
0
66
66.5
67
67.5
chrX (Mb)
RPKM
12.5
10.0
7.5
5.0
2.5
0.0
E1
E1b
E2
CE4
E3
CE1
CE2
CE3
E4
E5
E6
E7
E8
1
25
1
1
1
9
16
1
2
2
2
1

**Figure S3**

### Figure S3. Comparison of genomic alterations in disease states.

**(A)** Alteration status of paired samples from the same patients before and after treatment. Known prostate driver genes with alteration status in any of the included samples are shown.

**(B)** Comparison of rearrangement frequency in different disease states of mCRPC. The known prostate cancer driver genes that were located within 1 Mb to any SRB region are included.

**(C)** Intragenic deletion event leading to loss of ligand binding domain of *AR* in sample DTB-124-BL. *AR* gene boundary (green) and its enhancer (yellow) are shown; concave arcs, intra-chromosomal SV events; convex arcs, inter-chromosomal SV events. Copy number values represent 10 kB bins and have been tumor purity corrected. The expression values of all known *AR* exons, including both canonical and cryptic ones, are shown in the top panel. In the bottom panel, the number of reads covering the junction sites of two exons are indicated by weighted arcs.

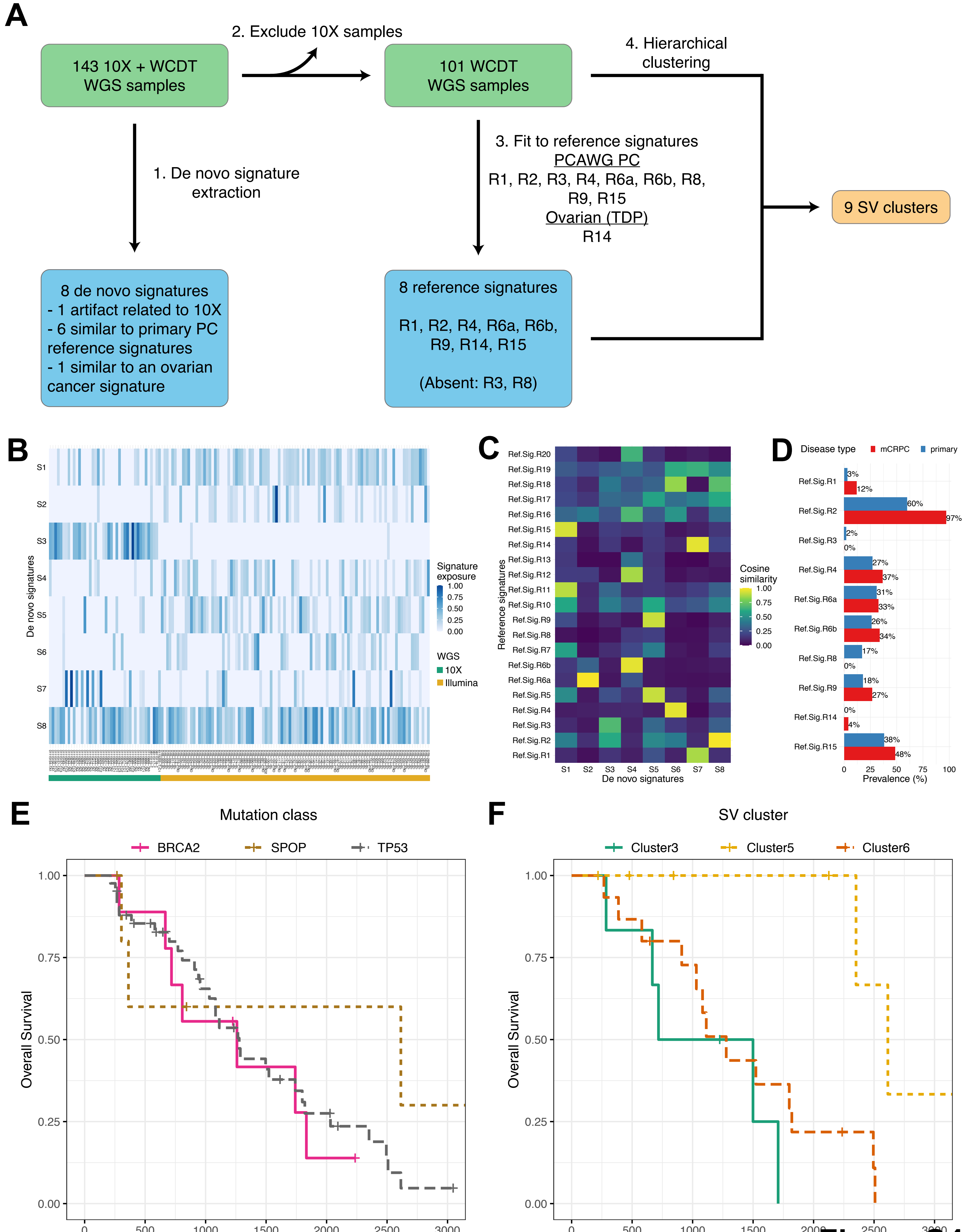
A
143 10X + WCDT WGS samples
2. Exclude 10X samples
101 WCDT WGS samples
4. Hierarchical clustering
1. De novo signature extraction
3. Fit to reference signatures
PCAWG PC
R1, R2, R3, R4, R6a, R6b, R8, R9, R15
Ovarian (TDP)
R14
9 SV clusters
8 de novo signatures
- 1 artifact related to 10X
- 6 similar to primary PC reference signatures
- 1 similar to an ovarian cancer signature
8 reference signatures
R1, R2, R4, R6a, R6b, R9, R14, R15
(Absent: R3, R8)
B
De novo signatures
S1
S2
S3
S4
S5
S6
S7
S8
Signature exposure
WGS
10X
Illumina
C
Reference signatures
Cosine similarity
De novo signatures
D
Disease type
mCRPC
primary
Prevalence (%)
E
Mutation class
BRCA2
SPOP
TP53
Overall Survival
Time (days)
F
SV cluster
Cluster3
Cluster5
Cluster6
Overall Survival
Time (days)

Figure S4

### Figure S4. Signature analysis of SV events.

**(A)** Workflow of SV signature analysis. Samples involved in this analysis are described in green boxes. Details of relevant signatures are shown in blue boxes. The steps for obtaining the final 9 SV clusters are indicated by numbers.

**(B)** Signature exposure based on de novo SV signatures. The exposure values of each sample were normalized such that the sample-wise sum is 1. Samples are ordered alphabetically based on names. The sequencing technology used for each sample is labeled at the bottom.

**(C)** Similarity between de novo SV signatures and reference signatures. Pairwise cosine similarity between de novo and reference SV signatures is shown.

**(D)** Comparison of reference signature (RefSig) prevalence between mCRPC and localized prostate cancer. The prevalence value for a signature in mCRPC was computed based on samples harboring at least 5% signature exposure. Localized prostate cancer prevalence values were obtained from signal.mutationalsignatures.com (48) computed from 199 PCAWG samples.

**(E)** Kaplan Meier curve of prediction using mutation class of key marker genes. Samples were grouped based on the mutation status of the corresponding marker gene.

**(F)** Kaplan Meier curve of prediction using SV cluster information. Samples were grouped based on their assignments of the corresponding SV cluster.

### SUPPLEMENTAL TABLE LEGENDS

#### Table S1. Sequencing, clinical and alteration information of all samples involved in this study.

**(A)** Sequencing metrics of all samples based on linked-read sequencing.

**(B)** Clinical properties and key genomics metrics of the cohort.

**(C)** Somatic mutation status of 159 prostate cancer drivers in the cohort. Sample and genes were sorted alphabetically. Genes with no detected mutations were left blank.

**(D)** Significantly mutated genes ($q \leq 0.1$) detected by dN/dS algorithm.

**(E)** Somatic copy-number alteration status of 159 prostate cancer drivers in the cohort.

**(F)** Recurrent copy-number alteration peaks detected by GISTIC.

**(G)** Gene transecting rearrangements of 159 prostate cancer drivers in the cohort. Types of rearrangement events were included.

**(H)** Gene flanking rearrangements of prostate cancer drivers in the cohort.

**(I)** TITAN copy number segments for all samples. Columns with "Corrected_*" were used for analysis in this study.

**(J)** TITAN optimal solutions selected for all samples.

**(K)** Structural variant calls for all samples. For samples with linked-read data ("CRPC10X"), union set of detected calls from SvABA, GROC-SVS, and Long Ranger are indicated. `SV.Filter` indicate SV events after filtering. `support` contain evidence from various callers; manual curation of events is indicated here. `CN_overlap_type` contain the final SV classification after annotation with copy number information.

**(L)** TDP status, copy number gain event counts, and median tandem duplication lengths for all samples.

#### Table S2. Significantly recurrent breakpoint regions and ETS fusion.

**(A)** Significantly recurrent breakpoints (SRB) regions ($q \leq 0.1$) in the mCRPC cohort of 143 samples.

**(B)** Significantly recurrent breakpoints (SRB) regions ($q \leq 0.1$) in the localized prostate cancer cohort of 278 samples.

**(C)** Fusion status of the ETS family genes. Gene expression was normalized to z-score for each gene. Genes with no detected fusion events or available expression data were left blank.

#### Table S3. *AR* alteration patterns in the mCRPC cohort.

#### Table S4. SV signature in the mCRPC cohort.

**(A)** Matrix of cosine similarity with rows representing reference signatures and columns representing *de novo* signatures.

**(B)** Exposure of all 8 *de novo* signatures in the cohort. Values were not normalized.

**(C)** Exposure of 8 chosen reference signatures in the cohort. Values were not normalized.
